# Genomic, effector protein and culture-based analysis of *Cyclaneusma minus* in New Zealand provides evidence for multiple morphotypes

**DOI:** 10.1101/2023.05.21.541640

**Authors:** M. Tarallo, K. Dobbie, L. Nunes Leite, T. Waters, K. Gillard, D. Sen, R.L. McDougal, C.H. Mesarich, R.E. Bradshaw

## Abstract

Cyclaneusma needle cast, caused by *Cyclaneusma minus*, affects *Pinus* species around the world. Previous studies suggested the presence of two distinct morphotypes in New Zealand, ‘verum’ and ‘simile’. Traditional mycological analyses revealed a third morphotype with clear differences in colony morphology and cardinal growth rates at varying temperatures. Genome sequencing of eight *C. minus* isolates provided further evidence of the existence of a third morphotype, named ‘novus’ in this study. To further analyse these morphotypes, we predicted candidate effector proteins for all eight isolates, and also characterized a cell-death eliciting effector family, Ecp32, which is present in other pine phytopathogens. In concordance with their distinct classification into three different morphotypes, the number of Ecp32 family members differed, with patterns of pseudogenization and some family members being found exclusively in some morphotypes. We also showed that proteins belonging to the Ecp32 family trigger cell death responses in non-host *Nicotiana* species, and, as previously demonstrated for other plant pathogens, the *C. minus* proteins belonging to the Ecp32 family adopt a β-trefoil fold. Understanding the geographical range and variations in virulence and pathogenicity of these morphotypes will provide a better understanding of pine needle diseases as well as enable the development of more durable methods to control this disease.

## Introduction

Cyclaneusma needle cast (CNC) is a disease of *Pinus* species in many parts of the world (Bednářová et al., 2013). CNC affects more than 30 pine species, including *Pinus radiata* in New Zealand and Australia, while in forests of Europe and North America, Scots pine (*Pinus sylvestris*) is prone to this disease (Merrill & Wenner, 1996; Müller & Kurkela, 2012). CNC results in premature needle loss, and therefore losses in growth and wood production. For New Zealand’s plantation forestry, the losses caused by CNC were estimated to be around $38 million per year in 2009 (Bulman, 2009), however this impact has likely now been reduced through the planting of less susceptible genotypes. Regardless, the disease persists in some locations, and there are no alternative control measures (Bulman, 1993).

Cyclaneusma needle cast is caused by the Leotiomycete fungus *Cyclaneusma minus* (Butin) DiCosmo, Peredo and Minter. At first, *C. minus* was reported as *Naemacyclus niveus* (Gilmour, 1959), which was later separated into two species, *N. niveus* and *Naemacyclus minor* (Von Butin, 1973). This separation was based on morphological characters on needles, the formation of the sexual and asexual stages in culture, and the relationship with different host species. *N. niveus* and *N. minor* were later reassigned to the genus *Cyclaneusma* as *Cyclaneusma niveum* and *C. minus*, respectively (Dicosmo et al., 1983). The morphological features distinguishing *C. niveum* and *C. minus* were described in Von Butin (1973; in German) and again in Dick et al. (2001; translated into English), where *C. niveum* isolates had sickle-shaped pycnidiospores and much shorter apothecia, and *C. minus* had bacilliform-shaped pycnidiospores and longer apothecia. Further investigation of the *C. minus* isolates in New Zealand found that, based on apothecial length variation as well as fungal morphology in culture, *C. minus* should be separated into two morphotypes: tagged as *C. minus* ‘verum’ and *C. minus* ‘simile’ (Dick et al., 2001). A relationship was shown between apothecium dimensions from needles and these morphotypes. A multigene phylogenetic analysis of New Zealand and Australian isolates of *Cyclaneusma* demonstrated that the separation of isolates by morphotype was consistent with genetic analysis and suggested the need for species descriptions to formally re-classify, as well as pathogenicity studies to understand phenotypic differences between the morphotypes (Prihatini et al., 2014).

Hunter et al. (2016) developed molecular tools for differentiating the ‘verum’ and ‘simile’ morphotypes and looked at their distribution throughout New Zealand. The study found that ‘simile’ was the most common morphotype in New Zealand, and that molecular classification was much more accurate than morphological assessments. Amongst the isolates studied, one (NZFS3325) showed DNA sequence variability in the ITS region of its genome, when compared to the corresponding region in both the ‘simile’ and ‘verum’, which indicated that additional morphotypes might exist (Hunter et al., 2016). Rightfully identifying these different morphotypes would help to determine their current distribution as well as to predict future distribution under different climatic scenarios, in New Zealand and globally. Comparative studies between morphotypes can also determine whether a morphotype is more virulent than others, all of which can have implications for disease diagnostics and forest management.

To further characterize the morphotypes of *C. minus*, we carried out genome sequencing for eight isolates: two belonging to morphotype ‘simile’ (Cms; NZFS759 and NZFS809), four belonging to morphotype ‘verum’ (Cmv; NZFS110, NZFS725, NZFS1800 and NZFS3276) and two isolates (Cmn; NZFS3305 and NZFS3325) that could not be differentiated into ‘verum’ or ‘simile’ using previously developed molecular tools. This genomic information will provide additional data to assist taxonomic classification.

An additional benefit of genomic resources is that they can provide insights into the molecular interactions of pathogens with their hosts, such as different *C. minus* morphotypes with *P. radiata*, through the prediction of candidate effector (CE) proteins. Effectors are usually small proteins secreted by pathogens into and around host cells to facilitate invasion, for example by suppressing host defence responses (Lo Presti et al., 2015; Rocafort et al., 2020). If the host has the ability to recognize such effectors, through its extracellular and intracellular resistance (R) proteins, immune responses are activated in order to stop pathogen growth (Cook et al., 2015; Jones & Dangl, 2006). This recognition can lead to a hypersensitive response (HR), which is generally characterized by a localized plant cell death (Win et al., 2012). Notably, pathogen effectors can be heterologously expressed in host and non-host plants, with those triggering a plant cell death response being candidates for recognition by a plant R protein.

In this study, genome sequencing was performed for eight *C. minus* isolates with the main objective of determining if a third morphotype is present in New Zealand. Moreover, the genome sequences of the different *C. minus* morphotypes were examined for genes encoding CE proteins, with a specific focus on homologues of the cell death-eliciting Ecp32 protein family, firstly identified in the tomato pathogen *Fulvia fulva* and subsequently in another pine pathogen, *Dothistroma septosporum* (Mesarich et al., 2018; Tarallo et al., 2022). Here, we compared their sequences, predicted tertiary structures, and screened these homologues for their ability to trigger cell death in the model non-host plants *Nicotiana benthamiana* and *Nicotiana tabacum* using *Agrobacterium tumefaciens*-mediated transient transformation assays (ATTAs). In doing so, another objective of this study was to provide novel insights into whether the Ecp32 family of *C. minus* plays an important role during host infection as well as to highlight potential differences in the molecular basis of how the different morphotypes interact with their host.

## Methods

### 1 Microorganisms and plants

*C. minus* isolates NZFS110, NZFS725, NZFS759, NZFS809, NZFS1800, NZFS3305, NZFS3276 and NZFS3325 (Table 1) were used in this study. Each was grown on a sterile cellophane sheet overlaying 2% (w/v) Difco^TM^ Malt Extract Agar (MEA) (Becton Dickinson & Company, New Jersey, USA) and incubated at 22°C in the dark. Once the mycelium filled half of the plate, it was scraped into a sterile 15 mL Falcon tube and stored at –80°C in preparation for freeze-drying and genomic DNA (gDNA) extraction.

**Table 1.**
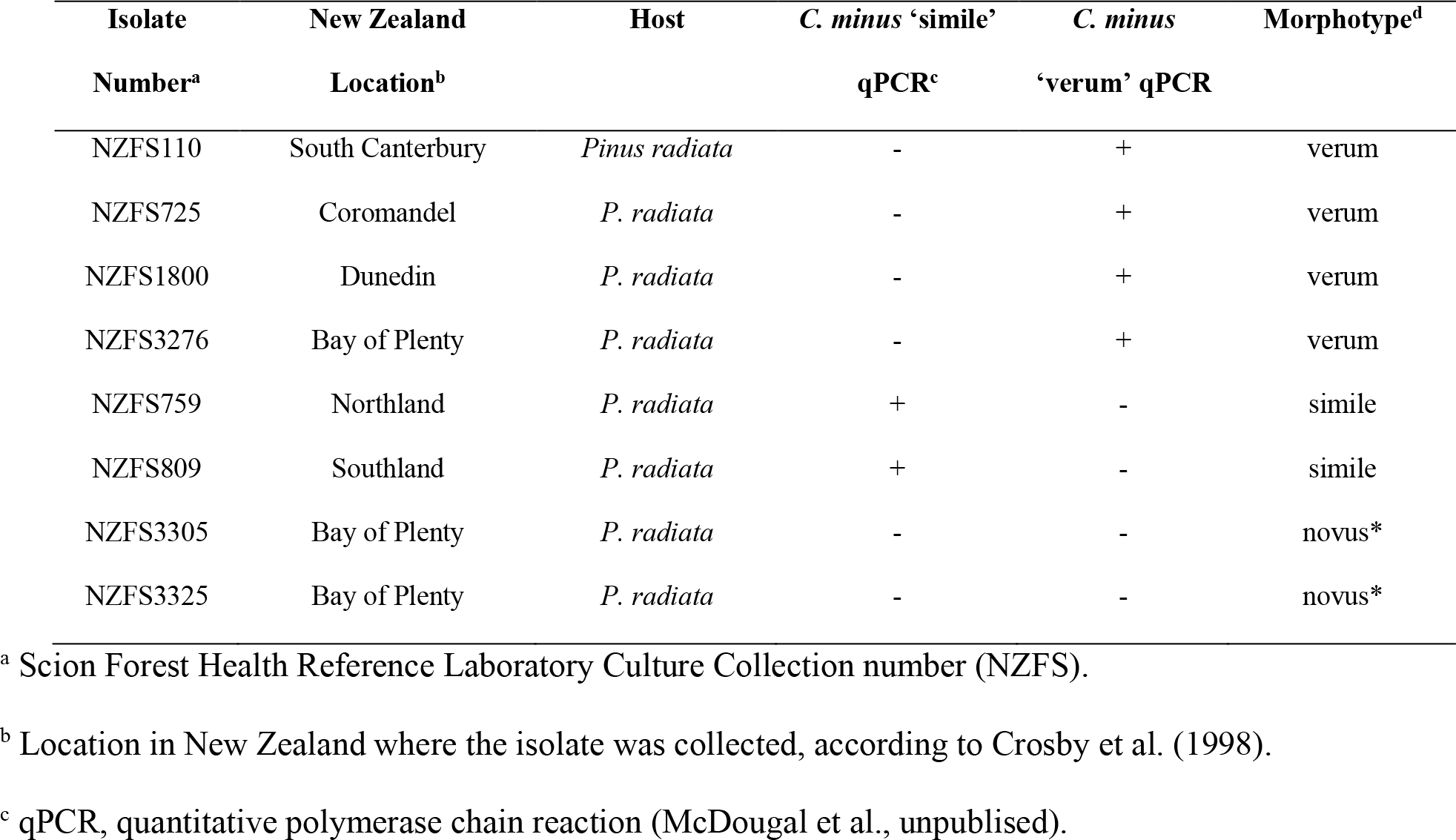
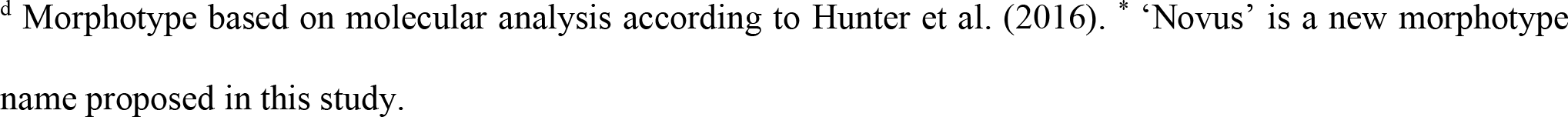
Isolates of *Cyclaneusma minus* used in this study.

| Isolate<br>Number<sup>a</sup> | New Zealand<br>Location<sup>b</sup> | Host | *C. minus* ‘simile’<br>qPCR<sup>c</sup> | *C. minus*<br>‘verum’ qPCR | Morphotype<sup>d</sup> |
| --- | --- | --- | --- | --- | --- |
| NZFS110 | South Canterbury | *Pinus radiata* | - | + | verum |
| NZFS725 | Coromandel | *P. radiata* | - | + | verum |
| NZFS1800 | Dunedin | *P. radiata* | - | + | verum |
| NZFS3276 | Bay of Plenty | *P. radiata* | - | + | verum |
| NZFS759 | Northland | *P. radiata* | + | - | simile |
| NZFS809 | Southland | *P. radiata* | + | - | simile |
| NZFS3305 | Bay of Plenty | *P. radiata* | - | - | novus* |
| NZFS3325 | Bay of Plenty | *P. radiata* | - | - | novus* |

<sup>a</sup> Scion Forest Health Reference Laboratory Culture Collection number (NZFS).
<sup>b</sup> Location in New Zealand where the isolate was collected, according to Crosby et al. (1998).
<sup>c</sup> qPCR, quantitative polymerase chain reaction (McDougal et al., unpublished).
<sup>d</sup> Morphotype based on molecular analysis according to Hunter et al. (2016). \* ‘Novus’ is a new morphotype name proposed in this study.

*Escherichia coli* DH5α (Taylor et al., 1993) was used for CE gene cloning, and *A. tumefaciens* GV3101 (Holsters et al., 1980) was used for ATTAs. *N. tabacum* Wisconsin 38 and *N. benthamiana* were used as model non-host plants for ATTAs. Plants were grown in a temperature-controlled room at 22°C and 12:12 light:dark photoperiod with 80-85 µmol m^-2^s^-1^.

### .2 Colony morphology analysis

For colony morphology assessments, three 5 mm diameter plugs from the edge of an actively growing culture of each *C. minus* isolate were placed in the middle of 2% and 9% MEA plates. Plates were incubated as above and assessed for colony morphology at eight weeks post-inoculation using a standard colour chart used to describe colours in colonies from Rayner (1970). For the temperature study, a 5 mm diameter plug from each isolate was placed onto an independent 2% MEA plate as before, each with three replicates, and incubated at 4, 10, 17, 20, 22, 25, 28 or 30°C in the dark. Two measurements at right angles for each replicate were taken at 14 days post-inoculation; the mean and standard deviation of the radial growth rate per day for each isolate were then calculated for each incubation temperature. The cardinal temperatures for each morphotype were determined from these data.

### .3 Genome sequencing, assembly, and annotation

Total gDNA for genome sequencing was extracted from freeze-dried mycelia of *C. minus* isolates using a DNeasy Plant Maxi Kit (Qiagen), according to the manufacturer’s instructions. For each isolate, 200 mg of tissue was added to Lysing Matrix C tubes containing 1.0 mm silica spheres (MP Biomedical, California, USA) and homogenised to a fine powder using a FastPrep-24^TM^ Instrument (MP Biomedical) with two cycles of 20 sec at 5 m/sec. The extracted gDNA was concentrated by ethanol precipitation and resuspended in TE buffer (10 mM Tris and 1 mM EDTA, pH 8). The gDNA concentrations were quantified by spectrophotometry using both the Denovix DS-11 FX (DeNovix, Delaware, USA) and the DeNovix dsDNA Broad Range Assay (Denovix).

The gDNA from eight isolates of *C. minus* (Table 1) was used for whole-genome sequencing using Illumina short-read sequencing technology (Illumina, California, USA). Paired-end libraries with insert sizes of 327–353 bp were sequenced on a NovaSeq 6000 system (Illumina).

*De novo* genome assemblies of filtered reads were generated using SPAdes v3.14.0 (Nurk et al., 2013) with default settings. Contigs were assembled into scaffolds with redundans v0.13c (Pryszcz & Gabaldón, 2016). Genome assembly quality was measured in QUAST (Gurevich et al., 2013). Assembly completeness was estimated with Benchmarking Universal Single-Copy Orthologs (BUSCO) comparisons to the fungal dataset (Simão et al., 2015).

Gene annotations were predicted in each masked genome assembly using the Funannotate v1.8.9 pipeline (Palmer & Stajich, 2020). Gene prediction was carried out using predicted protein sequences of *Elytroderma deformans* CBS183.68 (available at: https://mycocosm.jgi.doe.gov/Elyde1/Elyde1.home.html) as a guide. At first, protein sequences of *E. deformans* were mapped to the *C. minus* assembly file and the orthologs from *Cyclaneusma* were extracted. These orthologous sequences were used to train Augustus, followed by three rounds of iterative gene prediction. Genes were predicted by Augustus v3.3.3 (Stanke et al., 2004), GeneMark ES v4.68 (Ter-Hovhannisyan et al., 2008), snap 2006-07-28 (Korf, 2004) and GlimmerHMM v3.0.4 (Majoros et al., 2004), while transfer RNAs were predicted by tRNAscan-SE v2.0.9 (Chan et al., 2021). EVidenceModeler v1.1.1 (Haas et al., 2008) was used to generate consensus gene models. Putative functions of gene products were determined by searches to the MEROPS v12.0 (Rawlings et al., 2018), UniProt DB 2021_03 (The UniProt, 2021), dbCAN v10.0 (Yin et al., 2012), PFAM v34.0 (Mistry et al., 2021) and BUSCO v2.0 (Simão et al., 2015) databases.

### .4 Phylogenetic trees

The phylogenetic positions of the *C. minus* strains were assessed by constructing phylogenetic trees using concatenated nucleotide sequences from the Internal Transcribed Spacer (ITS) of ribosomal DNA, nuclear LSU (nLSU) and translation elongation factor 1α (tef-1). An initial alignment of these barcode genes from *Lophodermium conigenum* NZFS790 to the assembled genomes was done using LASTZ (Harris, 2007) and Geneious R10 (Kearse et al., 2012) to find the *C. minus* orthologs. Orthologs were then extracted and aligned to DNA sequences from other *Cyclaneusma* species, as well as *Lophodermium* and *Strasseria* species (Additional File 1). The alignment for the tree was generated using a cost-matrix of 51% with pairwise alignment or multiple alignment gap opening penalty of 12, plus a gap extension penalty of 0.3. The DNA sequences were aligned in Geneious R10. Nucleotide sequences not aligned at the 5’ and 3’ extremities of alignments were trimmed, and internal gaps maintained. The multi-gene phylogenetic trees were created by a maximum likelihood algorithm with 1000 replicates and distances obtained using the General Time Reversible model (GTR) using Mega X (Kumar et al., 2018).

### .5 Effector prediction

Following genome assembly and annotation of the eight *C. minus* isolates, a previously described pipeline (Hunziker et al., 2021) was used to identify CE proteins from all genomes, based on their full sets of predicted proteins. Briefly, signal peptides were predicted using SignalP v3.0 (Bendtsen et al., 2004). To exclude membrane-bound proteins, TMHMM v2.0 (Krogh et al., 2001) and BIG-PI (Eisenhaber et al., 2004) were used. Predicted proteins of ≤300 amino acids in length with ≥4 cysteine residues were then selected and classified as small, secreted and cysteine-rich proteins (SSCPs). EffectorP v3.0 (Sperschneider & Dodds, 2021) was then used to select proteins classified as apoplastic or cytoplasmic effectors, with candidates that were classified as non-effectors being discarded.

The amino acid sequence of DsEcp32-1, a member of the Ecp32 effector family from *D. septosporum* (Tarallo et al., 2022), was firstly used to identify the likely orthologue in *C. minus* isolate NZFS809. This *C. minus* sequence was then used to identify possible Ecp32 family homologues in the other seven isolates of *C. minus*, through reciprocal BLASTp and tBLASTn searches in conjunction with an E-value cut-off of <10^-5^.

Protein sequence alignments were performed using Clustal Ω (Sievers et al., 2011) in Jalview (Waterhouse et al., 2009). Phylogenetic trees were constructed with the neighbour-joining method based on 1000 bootstrap replicates using Geneious Software v9.1.8 (Kearse et al., 2012) in conjunction with CE protein sequence alignments.

Intrinsically disordered regions (IDRs) were predicted by querying protein sequences in the Predictor of Natural Disordered Regions (PONDR® VLXT) server (Romero et al., 2001). A prediction score ≥0.8 was considered a significant IDR prediction.

Protein tertiary structure predictions of Ecp32 family members were performed using AlphaFold2 in conjunction with ColabFold (Jumper et al., 2021; Mirdita et al., 2022). Two values were used to assess the confidence of the predictions: the predicted local-distance difference test (pLDDT) score, which ranges from 0 to 100, with values closer to 100 indicating a high confidence in prediction, and the global superposition metric template modelling score (TM-score), ranging from 0 to 1, with a value of ≥0.5 indicating a similar fold between structures. The Dali server (Holm, 2020) was used to identify proteins with structural similarity in the Research Collaboratory for Structural Bioinformatics Protein Data Bank (RCSB PDB). Here, a Dali Z-score of ≥2 was used to infer structural similarity. Protein tertiary structures were visualized and rendered in PyMOL v2.5, with alignments carried out using the CEalign tool (DeLano, 2002).

### 6 *Agrobacterium tumefaciens*-mediated transient expression assays (ATTAs)

*C. minus* ATTA expression vectors were generated using the method described by Guo et al. (2020). The expression vector used was pICH86988, which contains a *N. tabacum* N-terminal PR1α signal sequence for apoplast secretion followed by an N-terminal 3xFLAG tag for western blotting detection (Weber et al., 2011). CE gene sequences of *C. minus*, excluding the nucleotide sequence encoding the predicted signal peptide, were synthetized by Twist Bioscience (California, USA), and included BsaI recognition sites and 4 bp overhangs specific for Golden Gate modular cloning (Engler et al., 2009). CE genes were also cloned into a version of the vector that lacked the sequence encoding the PR1α signal peptide, but that contained only a start codon. Constructs were transformed into competent cells of *E. coli* cells by electroporation and plasmids were extracted using an E.Z.N.A.^®^ Plasmid DNA Mini Kit I (Omega Bio-tek, Georgia, USA). Correct assemblies were confirmed by sequencing in association with the Massey Genome Service (Palmerston North, New Zealand). Expression vectors were then transformed into competent cells of *A. tumefaciens* by electroporation (Guo et al., 2020).

To perform the ATTAs, transformed cells of *A. tumefaciens* containing the CE genes were inoculated into lysogeny broth (LB) supplemented with 50 µg/mL kanamycin, 10 µg/mL rifampicin and 30 µg/mL gentamycin and grown overnight at 28°C in the dark until an OD600 of 0.4–2.0 was reached. Cells were collected by centrifugation at 2500 x *g* for 5 min and resuspended in 1 mL of infiltration buffer (10 mM MgCl2:6H2O, 10 mM MES-KOH, 100 μM acetosyringone (Sigma-Aldrich, Missouri, USA)). The OD600 was measured again, and sterile water added to reach a final OD600 of 0.5. These cultures were then incubated for 3 h at room temperature before infiltration. An *A. tumefaciens* transformant carrying an expression vector for the extracellular elicitin INF1, from *Phytophthora infestans* (Kamoun et al., 1997), was used as a positive control and the empty pICH86988 vector was used as a negative control. The abaxial surface of *N. benthamiana* and *N. tabacum* leaves was infiltrated with the appropriate *A. tumefaciens* solution using 1 mL needleless syringes. The screening was done with at least three biological replicates. Each biological replicate consisted of two plants, each with two leaves infiltrated, resulting in at least 12 infiltration zones. Plants were monitored for up to seven days, at which time photographs were taken with a Nikon D7000 camera. To assess the plant response caused by each expressed protein, infiltration zones were divided into a strong or weak cell death response, chlorosis, or no cell death. A strong cell death response was considered when it was indistinguishable from the response elicited by INF1 and covered the entire infiltration zone, while a weak response was considered when the protein elicited a response across approximately 50% of the infiltration zone when compared with INF1. Chlorosis was considered when leaf discoloration occurred, but no apparent cell death was observed. A no-cell death response was indicated when the infiltration zone had the same lack of a response as the negative empty pICH86988 vector control.

## Results

### 1 *Cyclaneusma minus* isolates can be differentiated into three morphotypes based on molecular and morphological analyses

Eight *C. minus* isolates were collected from different parts of New Zealand and classified into two different morphotypes, ‘verum’ (Cmv) and ‘simile’ (Cms), according to molecular analysis (Table 1, Additional File 2) (Hunter et al., 2016). However, two of these isolates, NZFS3305 and NZFS3325 (NZFS3325 was previously identified in Hunter et al. (2016)), could not be differentiated based on this molecular analysis and were, therefore, classified as a new *C. minus* morphotype, ‘novus’ (Cmn). The phylogeny suggested that morphotype Cmn is more closely related to Cms than to Cmv (Additional File 2).

This classification was also supported by morphological analyses. Isolates and their corresponding morphotypes grown on 2% and 9% MEA at 22°C in the dark after 8 weeks are shown in Figure 1. Additional File 3 describes the colony morphology of each morphotype on 2 % and 9% MEA at 22 °C in the dark after 8 weeks. Results for the temperature study are shown in Additional File 4. The widest temperature range for growth, as well as the fastest growth rate, was seen in the Cmv morphotype. It had an optimal growth at 25°C, with a mean radial growth rate of 2.0 ± 0.7 mm day^-1^ at 25°C. However, growth at our lowest (4°C) and highest (30°C) temperatures were also detected for this morphotype. The Cms and Cmn morphotypes shared similar optimal growth temperatures at 20°C and 22°C, respectively, while growth ceased at 4°C and 30°C for both. The Cms morphotype had a slightly faster mean radial growth rate of 0.96 ± 0.13 mm day^-1^ at its optimal growth temperature of 20°C, compared to that of the Cmn morphotype being 0.69 ± 0.15 mm day^-1^ at 22°C.

**Figure 1.**
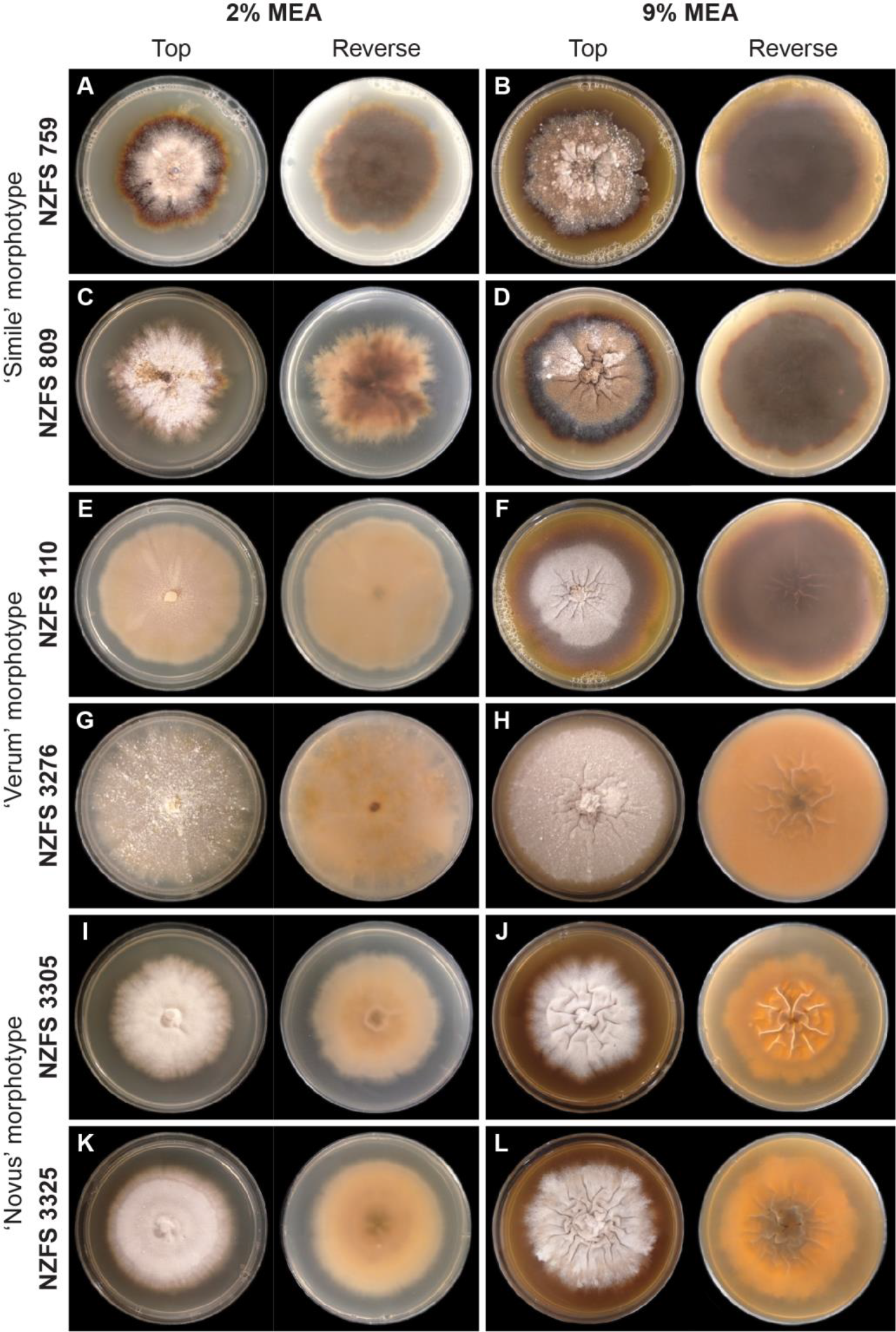
Colony morphology of *Cyclaneusma minus* isolates supports their classification as different morphotypes. All isolates were grown at 22°C in the dark for 8 weeks.

### 2 Differences in *Cyclaneusma minus* morphotype are supported by differences in genome size

The final genome assemblies for the eight *C. minus* isolates ranged in length from 31 to 43 Mb (Table 2 and Additional File 5). The final set of gene models for each genome showed a completeness of >96% with the retrieval of at least 728 complete BUSCO hits from a total of 758 fungal BUSCOs (Table 2 and Additional File 5). The Cmv morphotype was predicted to have a larger genome size (based on assembly length) and higher number of gene models compared to the Cms and Cmn morphotypes (Table 2).

**Table 2.**
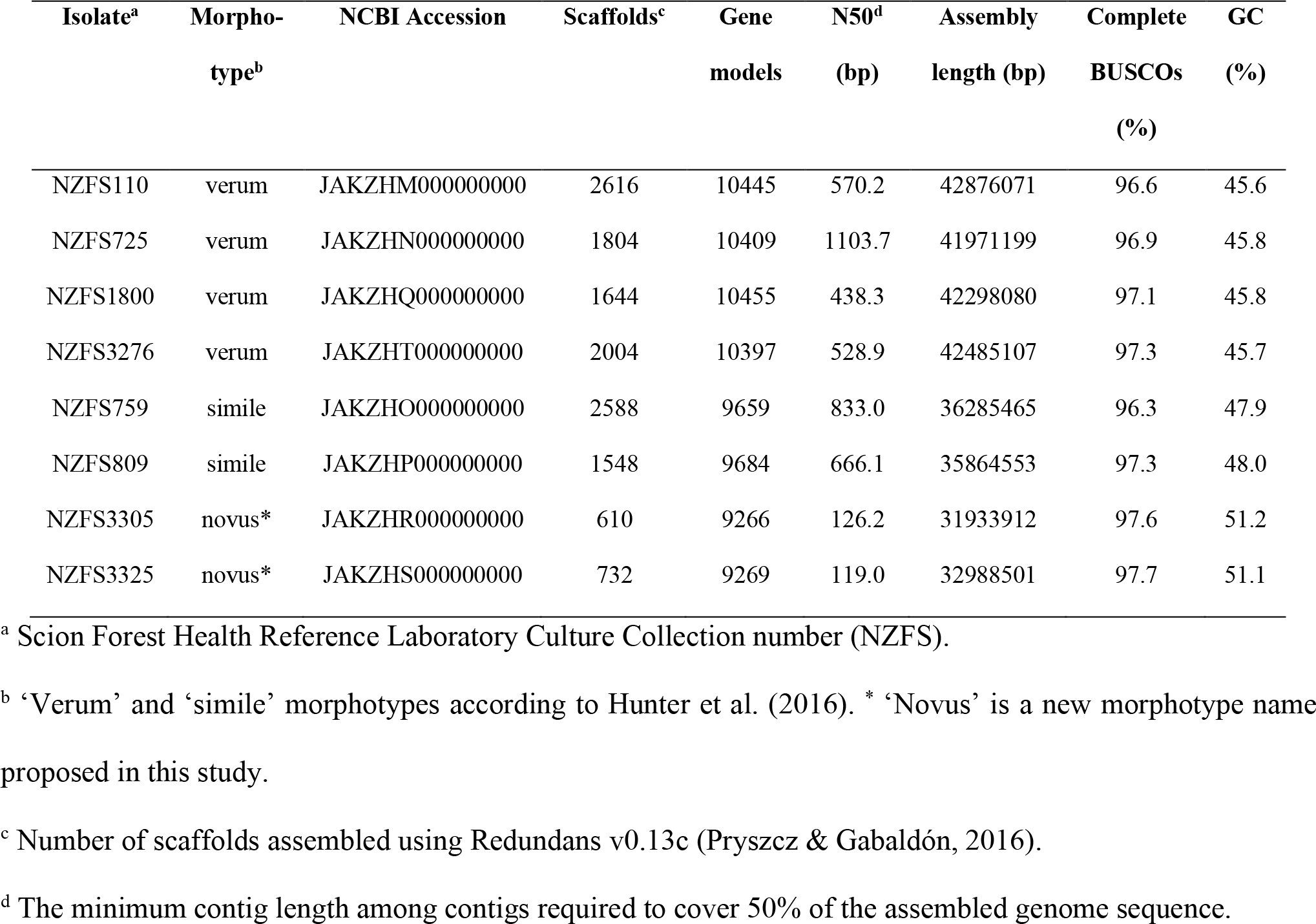
*Cyclaneusma minus* genomes statistics.

| Isolate<sup>a</sup> | Morpho-<br>type<sup>b</sup> | NCBI Accession | Scaffolds<sup>c</sup> | Gene<br>models | N50<sup>d</sup><br>(bp) | Assembly<br>length (bp) | Complete<br>BUSCOs<br>(%) | GC<br>(%) |
| --- | --- | --- | --- | --- | --- | --- | --- | --- |
| NZFS110 | verum | JAKZHM000000000 | 2616 | 10445 | 570.2 | 42876071 | 96.6 | 45.6 |
| NZFS725 | verum | JAKZHN000000000 | 1804 | 10409 | 1103.7 | 41971199 | 96.9 | 45.8 |
| NZFS1800 | verum | JAKZHQ000000000 | 1644 | 10455 | 438.3 | 42298080 | 97.1 | 45.8 |
| NZFS3276 | verum | JAKZHT000000000 | 2004 | 10397 | 528.9 | 42485107 | 97.3 | 45.7 |
| NZFS759 | simile | JAKZHO000000000 | 2588 | 9659 | 833.0 | 36285465 | 96.3 | 47.9 |
| NZFS809 | simile | JAKZHP000000000 | 1548 | 9684 | 666.1 | 35864553 | 97.3 | 48.0 |
| NZFS3305 | novus* | JAKZHR000000000 | 610 | 9266 | 126.2 | 31933912 | 97.6 | 51.2 |
| NZFS3325 | novus* | JAKZHS000000000 | 732 | 9269 | 119.0 | 32988501 | 97.7 | 51.1 |

<sup>a</sup> Scion Forest Health Reference Laboratory Culture Collection number (NZFS).
<sup>b</sup> ‘Verum’ and ‘simile’ morphotypes according to Hunter et al. (2016). \* ‘Novus’ is a new morphotype name proposed in this study.
<sup>c</sup> Number of scaffolds assembled using Redundans v0.13c (Pryszcz & Gabaldón, 2016).
<sup>d</sup> The minimum contig length among contigs required to cover 50% of the assembled genome sequence.

### 3 Prediction of candidate effector proteins from *Cyclaneusma minus* morphotypes

Effectors can have virulence functions during host infection by supressing plant defence responses or can be avirulence factors, in which case they are recognized by R proteins from the host. For this reason, following genome sequencing and assembly, secreted CE proteins were identified from the predicted protein set of all eight *C. minus* isolates to determine whether their CE protein repertoires were morphotype-specific. Strikingly, genomes from the same morphotype were predicted to have similar numbers of proteins and effectors, with Cmv having more predicted CEs than Cms or Cmn (Table 3). In all cases, a higher number of apoplastic effectors, when compared to cytoplasmic effectors, were predicted in all isolates analysed. Additional File 6 shows all amino acid sequences of the CEs predicted from the isolates with EffectorP.

**Table 3.**
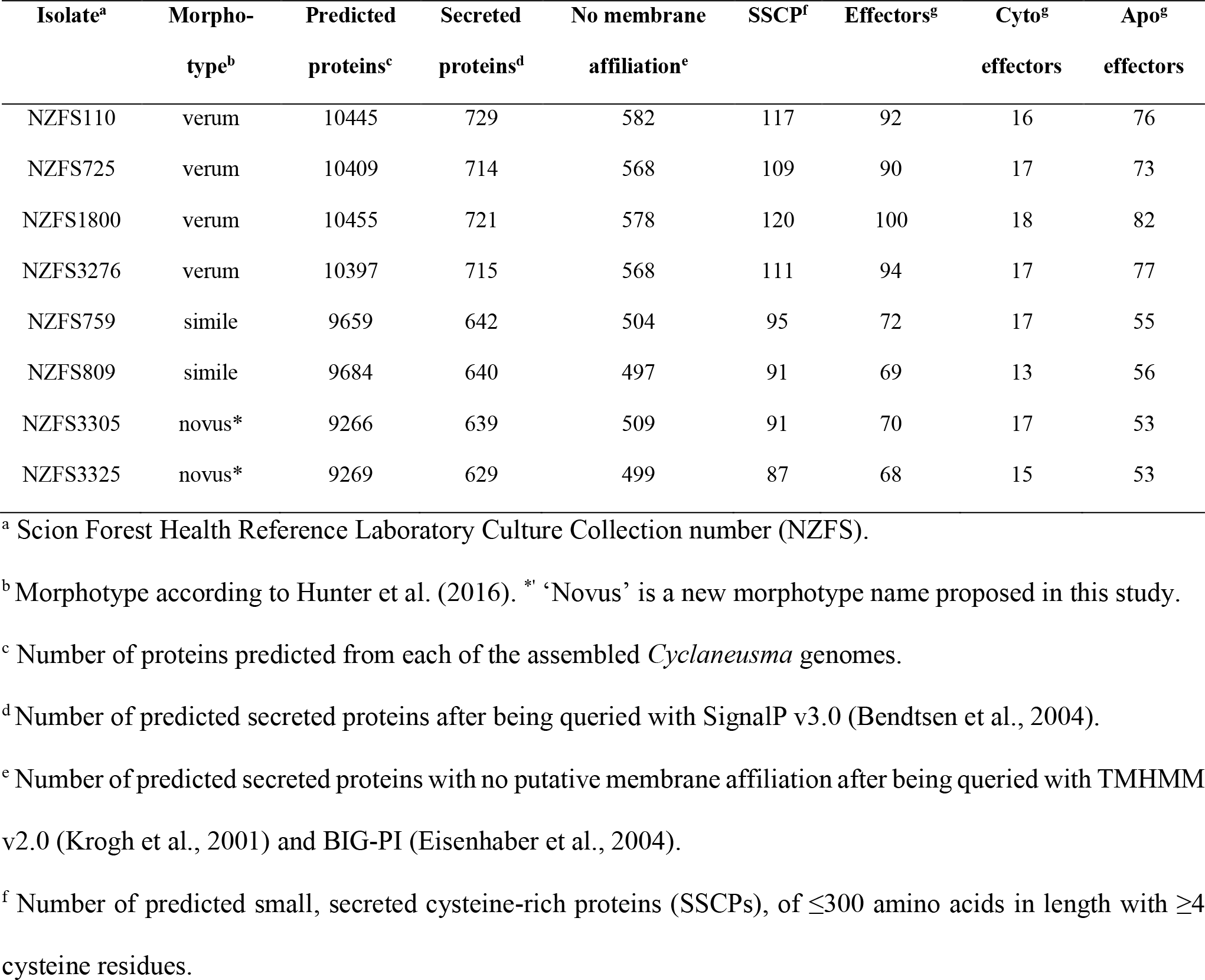
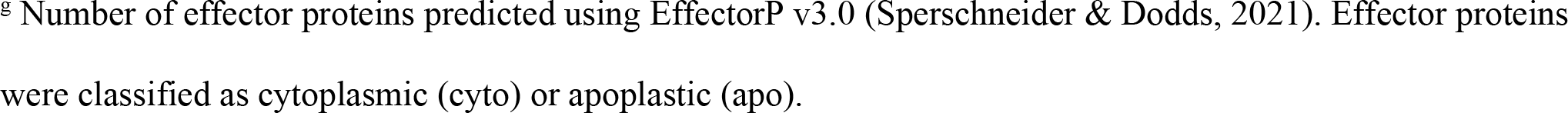
Effector protein prediction statistics across the eight *Cyclaneusma minus* isolates investigated in this study.

| Isolate<sup>a</sup> | Morpho-<br>type<sup>b</sup> | Predicted<br>proteins<sup>c</sup> | Secreted<br>proteins<sup>d</sup> | No membrane<br>affiliation<sup>e</sup> | SSCP<sup>f</sup> | Effectors<sup>g</sup> | Cyto<sup>g</sup><br>effectors | Apo<sup>g</sup><br>effectors |
| --- | --- | --- | --- | --- | --- | --- | --- | --- |
| NZFS110 | verum | 10445 | 729 | 582 | 117 | 92 | 16 | 76 |
| NZFS725 | verum | 10409 | 714 | 568 | 109 | 90 | 17 | 73 |
| NZFS1800 | verum | 10455 | 721 | 578 | 120 | 100 | 18 | 82 |
| NZFS3276 | verum | 10397 | 715 | 568 | 111 | 94 | 17 | 77 |
| NZFS759 | simile | 9659 | 642 | 504 | 95 | 72 | 17 | 55 |
| NZFS809 | simile | 9684 | 640 | 497 | 91 | 69 | 13 | 56 |
| NZFS3305 | novus* | 9266 | 639 | 509 | 91 | 70 | 17 | 53 |
| NZFS3325 | novus* | 9269 | 629 | 499 | 87 | 68 | 15 | 53 |

<sup>a</sup> Scion Forest Health Reference Laboratory Culture Collection number (NZFS).
<sup>b</sup> Morphotype according to Hunter et al. (2016). \* ‘Novus’ is a new morphotype name proposed in this study.
<sup>c</sup> Number of proteins predicted from each of the assembled *Cyclaneusma* genomes.
<sup>d</sup> Number of predicted secreted proteins after being queried with SignalP v3.0 (Bendtsen et al., 2004).
<sup>e</sup> Number of predicted secreted proteins with no putative membrane affiliation after being queried with TMHMM v2.0 (Krogh et al., 2001) and BIG-PI (Eisenhaber et al., 2004).
<sup>f</sup> Number of predicted small, secreted cysteine-rich proteins (SSCPs), of $\leq 300$ amino acids in length with $\geq 4$ cysteine residues.
<sup>§</sup> Number of effector proteins predicted using EffectorP v3.0 (Sperschneider & Dodds, 2021). Effector proteins were classified as cytoplasmic (cyto) or apoplastic (apo).

### 4 The number of Ecp32 effector family members differs between the three morphotypes of *Cyclaneusma minus,* further supporting their differentiation

To investigate potential differences in the effector protein repertoires of *C. minus* morphotypes, we focused on the Ecp32 effector family that had been characterised in another fungal pine pathogen (Tarallo et al., 2022). An apoplastic protein from isolate NZFS809 (Cms), Cms835, was firstly identified as an orthologue of DsEcp32-1 from *D. septosporum* and shown to be part of the Ecp32 effector family. With the Cms835 (NZFS809) amino acid sequence as reference, other Ecp32 family members in this and other isolates/morphotypes were identified from their respective proteomes and genomes using BLASTp and tBLASTn, respectively. Based on this analysis, it was determined that all isolates of all morphotypes had at least three members of the Ecp32 family of proteins (Figure 2 and Additional File 7).

**Figure 2.**
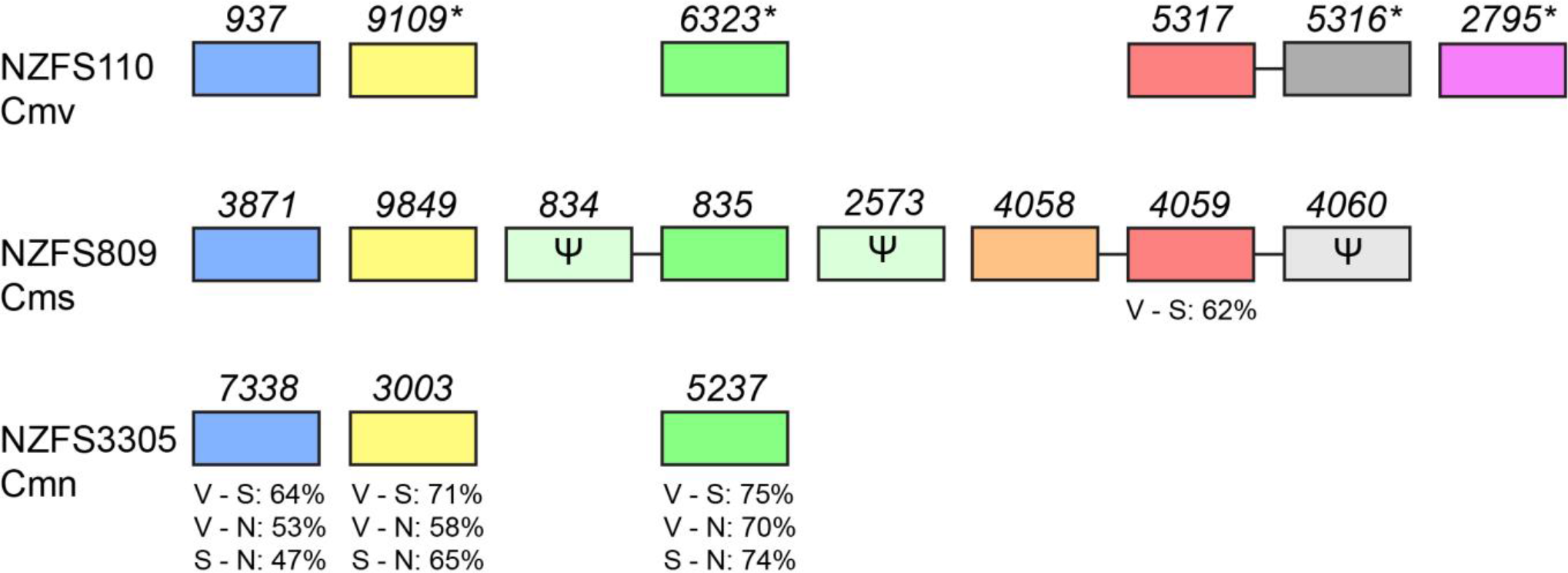
Schematic of the *Ecp32* family from three different morphotypes of *Cyclaneusma minus*. One representative isolate from each morphotype was chosen: NZFS110 for morphotype ‘verum’ (Cmv), NZFS809 for morphotype ‘simile’ (Cms), and NZFS3305 for morphotype ‘novus’ (Cmn). *Cms835* (green) was the first *Ecp32* family member identified, based on similarity with *DsEcp32-1* from the pine pathogen *Dothistroma septosporum*. Each rectangle represents a gene and orthologous genes between morphotypes are colour-coded in the same way as Figure 3. Ψ refers to pseudogenes. The horizontal lines connecting the rectangles indicate the genes are on the same scaffold and adjacent in the genome. The percentages refer to full-length pairwise amino acid identity between orthologous proteins encoded by each gene from each representative morphotype (V = ‘verum’, S = ‘simile’, N = ‘novus’). The pink rectangle indicates the gene encoding the protein with a predicted intrinsically disordered region. An asterisk * next to the gene number indicates amino acid variants in the respective encoded proteins that occurred in one or more of the other Cmv isolates. NZFS3276 orthologues of 9109, 6323 and 2795 have variants I34T, V188I and E40D, respectively. Orthologues of the other polymorphic protein indicated, 5316, have a Q82H substitution in all three other Cmv isolates and a S215G substitution in isolates NZFS1800 and 3276.

Even though all isolates and morphotypes appeared to have proteins belonging to the Ecp32 family, the number of family members between each morphotype varied. A BLASTp search indicated that the NZFS759 and NZFS809 genomes (both Cms) each encode eight Ecp32 family proteins. Each of the Cmv isolates (NZFS110, NZFS725, NZFS1800 and NZFS3276) have six paralogues in their Ecp32 family, while Cmn isolates (NZFS3305 and NZFS3325) have only three. The *Ecp32* genes from one representative isolate of each morphotype are illustrated in Figure 2. Unlike what was observed for orthologues between *D. septosporum* and *F. fulva*, proteins from the Ecp32 family from each *C. minus* isolate did not group with the previously identified members of the Ds/FfEcp32 family (Additional File 8). As was also found for the *D. septosporum* and *F. fulva* Ecp32 proteins, all Ecp32 family members from *C. minus* have predicted signal peptides but no known functional domains.

Some Ecp32 family members are found exclusively in one morphotype. This is the case for a set of orthologous proteins (Cmv2795, Cmv13188, Cmv116 and Cmv3437) from the four Cmv isolates. These proteins have longer amino acid sequences than the rest of the proteins in this family. Normally they range from 197 to 226 amino acids in length, however these four proteins each contain 268 amino acids; the additional amino acids form a predicted intrinsically disordered region (IDR) at the N-terminus, immediately following the putative signal peptide (Additional File 9).

The morphotypes also showed differences in pseudogenisation of *Ecp32* genes. For Cms isolates NZFS809 and NZFS759, three of the eight *Ecp32* genes were pseudogenes that were predicted to encode truncated proteins, with identical mutations in the homologues from these two isolates (NZFS809 is shown in Figure 2). Two of the genes (*Cms834* and *Cms4060*) had a premature stop codon at the position encoding amino acids 54 and 149, respectively, whilst *Cms2573* had a cytosine insertion at nt position 213 in the coding sequence, causing a frameshift that also led to a premature stop codon. Interestingly, in both Cms isolates, *Cms834* (and the NZFS759 orthologue) is on the same scaffold and adjacent to *Cms835* (and the NZFS759 orthologue) (Figure 2). Similarly, another gene cluster in this morphotype includes the *Cms4060* pseudogene, alongside *Cms4058* and *Cms4059*, which are predicted to encode functional proteins (Figure 2). This is also true for the respective orthologues from NZFS759.

Given that the Cms and Cmn morphotypes are the most closely-related of the three morphotypes based on the multigene phylogeny generated in this study (Additional File 2), the difference in the number of *Ecp32* gene family members between them was striking, even though three of the eight Cms genes were pseudogenised (Figure 2). Furthermore, the three genes that were shared orthologues between Cms and Cmn showed sequence diversity, with predicted amino acid sequence identities of only 47%, 65% and 74% between these morphotypes (Figure 2). Together, these results support the identification of Cmn as a distinct morphotype from Cms and Cmv, and also suggest that several gene duplication events and possibly gene loss may have occurred in this effector family, either in *C. minus* or in an ancestral species.

### 5 The Ecp32 family from different *Cyclaneusma minus* morphotypes are sequence-divergent and some members are found exclusively in certain morphotypes

To further investigate differences between the sets of Ecp32 family members across the morphotypes and isolates of *C. minus*, we performed a protein sequence alignment of all full-length members from the Ecp32 family (Additional File 9). Family members that had identical amino acid sequences were excluded from this alignment, as were sequences encoded by pseudogenes. Figure 3 shows a phylogenetic tree constructed based on these alignments, including all family members identified, even those with identical amino acid sequences, but still excluding those encoded by pseudogenes.

**Figure 3.**
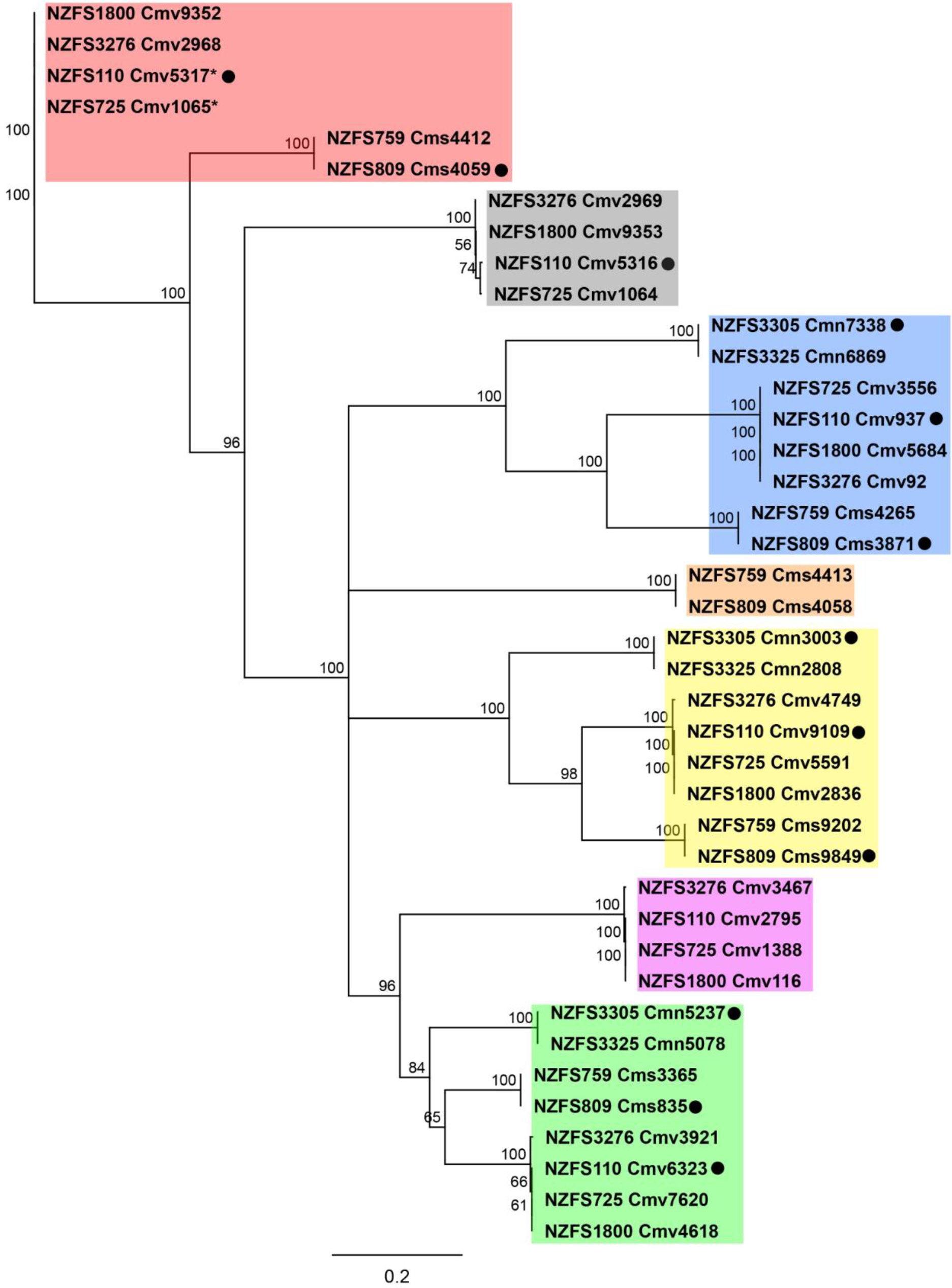
Phylogenetic tree of *Cyclanuesma minus* Ecp32 family members across eight isolates and three morphotyes of *Cyclanuesma minus*. Neighbour-joining tree obtained from the amino acid sequence alignment of Ecp32 proteins using Geneious Software v9.1.8. Bootstrap values are shown at the nodes as percentages. The scale bar represents 0.2 substitutions per site. The number following each morphotype ID (NZFS) refers to the protein number from the annotated genomes; the asterisk * indicates that the protein was incorrectly predicted, and the corrected protein sequence was used. The sequences encoded by predicted pseudogenes were excluded from the analysis. The black dot refers to the proteins that were expressed in *Nicotiana benthamiana* and *Nicotiana tabacum* using an *Agrobacterium tumefaciens*-mediated transient expression assay to assess their ability to elicit cell death. Cms: *C. minus* morphotype ‘simile’; Cmv: *C. minus* morphotype ‘verum’; Cmn: *C. minus* morphotype ‘novus’. Proteins were colour-coded according to their clustering in the phylogeny.

The results affirm that proteins from the Ecp32 family are more expanded in some morphotypes than in others, and they cluster into seven separate groups (colour-coded to match Figure 2). The clustered groups coloured in blue, yellow and green are made up of eight protein members, with one paralogue belonging to each of the eight genomes analysed. The clustered group colour-coded in red has six members, with paralogues only belonging to Cms and Cmv, and no members belonging to Cmn. The pink group corresponds to Cmv proteins with a predicted disordered region of 73 additional amino acids, which are most closely related to green group proteins, despite their larger size.

A high level of sequence divergence was seen between the Ecp32 proteins from the three morphotypes. Between all orthologous Cms and Cmv Ecp32 protein family members, the pairwise amino acid identity varied between 25.1% and 71.4%. Likewise, between Cms and Cmn, the pairwise identity was between 32.1% and 68.9%, and between Cmv and Cmn it was 28.8% and 68.6%. Differences were also observed between isolates of the same morphotype, as the amino acid sequence identity among the Ecp32 proteins ranged from 25.3% to 50.6% (Cmv), 30.9% to 42.3% (Cms), and 34.4% to 46.8% (Cmn).

We next compared the locations of clustered *Ecp32* genes with their positions in the phylogeny. All genes encoding Ecp32 proteins of the red and grey groups in the Cmv morphotypes are adjacent to each other in their respective genomes, as shown for *Cmv5316* and *Cmv5317* of isolate NZFS110 in Figure 2; the corresponding proteins are more similar in sequence to each other than to any other Ecp32 protein from that isolate, with identities ranging from 26.8% to 50.6% (between *Cmv5316* and *Cmv5317*) (Additional File 9). Together, these results suggest that these ‘red and grey’ family members may have arisen by an ancestral tandem duplication event. A similar linkage of these two family members was seen in the Cms morphotype (genes *Cms4059* and *Cms4060*), although in that case the ‘grey’ *Cm*s4060 gene was pseudogenised (Figure 2). Although not shown in the tree, Cms4060 showed greatest similarity with the proteins colour-coded in grey (Additional File 9), which suggests a common ancestral origin for the red and grey-coloured gene pairs of the Cmv and Cms morphotypes. A third *Ecp32* gene was also present in the red-grey gene cluster in the Cms morphotype: *Cms4058* (orange) (Figure 2). However, despite its close proximity to *Cms4059* in the genome, Cms4058 was more similar in sequence to Cms835 (green) than to Cms4059 (red).

Together, these results highlight the expansion of the *Ecp32* family in the three different morphotypes analysed, with possible gene duplication, sometimes in tandem, generating new family members. Sequence diversification was observed, not only between morphotypes, but also between family members from the same isolate.

### 6 Proteins from the Ecp32 family from different *Cyclaneusma minus* morphotypes trigger cell death in *Nicotiana* species

The next questions we addressed were whether the Ecp32 family proteins identified from the *C. minus* morphotypes could trigger cell death in *Nicotiana* (model non-host) species, similar to what was observed in *D. septosporum* and *F. fulva* (Tarallo et al., 2022), and whether there were differences in cell death-eliciting ability between morphotypes. The selected proteins tested by ATTAs are indicated with a black dot in Figure 3. One protein from one representative isolate of each morphotype was selected from the clusters that had a higher number of family members.

In *N. benthamiana*, all proteins tested triggered some degree of cell death, except for Cmv937 (the Cmv representative from the blue cluster in Figure 3) (Figure 4). Among those tested, proteins that consistently triggered cell death were those from the green cluster shown in Figure 3 (Cmv6323, Cms835 and Cmn5237), and the yellow cluster (Cmv9109, Cms9849 and Cmn3003), with Cms9849 triggering a strong cell death response in all infiltration zones. Across all proteins tested, differences were seen between morphotypes (Figure 4). The truncated protein Cms4060 (NZFS809) was also tested and triggered a weak cell death response. Although this protein was not included in the phylogenetic tree shown in Figure 3, it showed greatest similarity with the proteins colour-coded in grey, and an alignment between these proteins (Additional File 9) indicated conservation of the N-terminus (post signal peptide), which could suggest that this region of the protein might be responsible for cell death elicitation, since Cmv5316 also triggered a cell death response (Figure 4).

**Figure 4.**
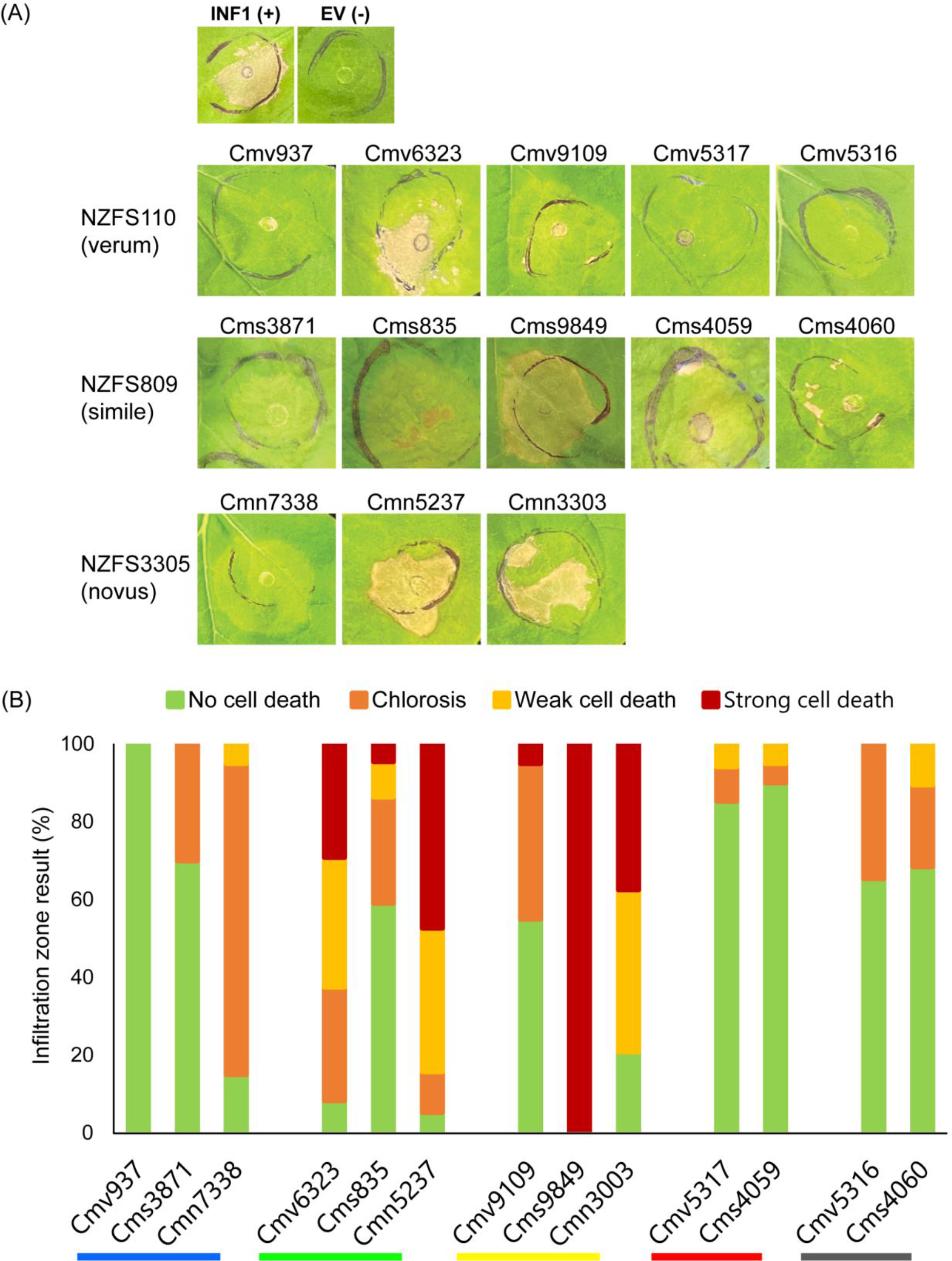
Ecp32 proteins from different morphotypes of *Cyclaneusma minus* trigger cell death in *Nicotiana benthamiana*. (A) Proteins were expressed in *N. benthamiana* using an *Agrobacterium tumefaciens*-mediated transient expression assay to assess their ability to elicit cell death. Representative images are shown (n = 12– 30 infiltration zones) from at least three independent experiments. INF1, *Phytophthora infestans* elicitin positive cell death control; EV, empty vector negative no-cell death control; Cms: *C. minus* morphotype ‘simile’; Cmv: *C. minus* morphotype ‘verum’; Cmn: *C. minus* morphotype ‘novus’. Photos were taken 7 days after infiltration. (B) Graphs display the percentages of infiltration zones that showed cell death in response to each different protein, divided into four categories: strong cell death, weak cell death, chlorosis, and no cell death. The proteins are shown as orthologous groups, with the different morphotypes represented for each. The coloured lines under the protein IDs refer to the different gene and protein groups in Figures 2 and 3.

In *N. tabacum*, yellow cluster proteins Cmv9109, Cms9849 and Cmn3003 (Figure 3) again consistently triggered strong cell death responses, while all other proteins inconsistently triggered either weak cell death or chlorosis (Additional File 10). Interestingly, the truncated protein Cms4060 (NZFS809), as well as its paralogue Cmv5316, did not trigger cell death (Additional File 10).

Taken together, these results suggest that different Ecp32 family members from the same isolate differ in their ability to trigger a cell death response in non-host plants. There were also differences in the responses observed for orthologues from different morphotypes (e.g., proteins from the blue cluster).

### 7 Predicted tertiary structures of the *Cyclaneusma minus* Ecp32 family members suggest possible roles in virulence

The tertiary structure of one *C. minus* Ecp32 family member from each of the clusters shown in Figure 3 was predicted using AlphaFold2 (Figure 5). The amino acid sequences of proteins from NZFS110 (Cmv) were used for these structural predictions since one Ecp32 family member from this isolate was present in all clusters, except for the orange one (Figure 3). All of the structures had high pLDDT scores and TM-scores, indicative of highly confident predictions (Additional File 11). As previously demonstrated for the Ecp32 proteins of *D. septosporum* and *F. fulva*, the Ecp32 family members from *C. minus* were predicted to be structurally similar to β-trefoil proteins, even though they had some differences in topology. The structures of Cmv937, Cmv6323, Cmv9109, Cmv5317 and Cmv2795 have two predicted disulphide bonds, while Cmv5316 has three.

**Figure 5.**
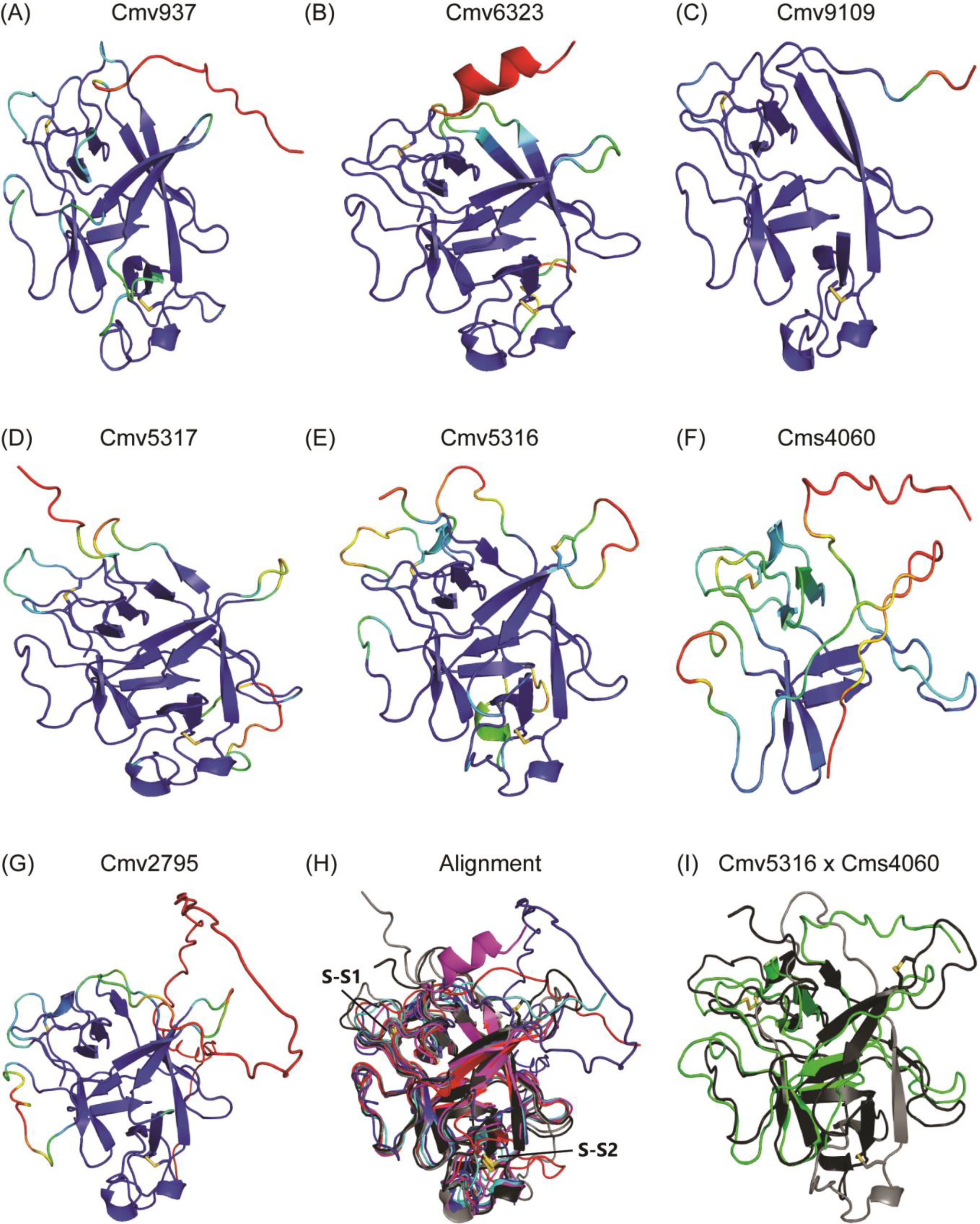
Predicted protein tertiary structures of Ecp32 family members from different isolates of *Cyclaneusma minus*. (A–G) Predicted structures of (A) Cmv937, (B) Cmv6323, (C) Cmv9109, (D) Cmv5317, (E) Cmv5316, (F) Cms4060 and (G) Cmv2795. Protein structures were predicted with AlphaFold2, rendered in PyMOL v2.5 (DeLano, 2002; Jumper et al., 2021; Mirdita et al., 2022), and coloured according to their AlphaFold2 pLDDT score: dark blue for regions predicted with high confidence, light blue and green for regions of low confidence, and red for very low confidence regions. (H) Alignment of the predicted Cmv937 (red), Cmv6323 (magenta), Cmv9109 (cyan), Cmv5317 (grey) and Cmv5316 (black) structures. S-S: disulphide bonds. (I) Alignment of the predicted Cmv5316 (black and grey) with the truncated Cms4060 (green). Black represents the part of the Cmv5316 structure that aligns with Cms460. Protein structures were aligned using the CEalign tool in PyMOL. Disulphide bonds are shown as yellow sticks. Cmv: *C. minus* morphotype ‘verum’; Cms: *C. minus* morphotype ‘simile’.

Pairwise structural alignments between all of the predicted structures suggested that Cmv6323 (green group) and Cmv9109 (yellow group) are more similar to one another than any other structural pairwise alignment from the predicted structures for Cmv isolate NZFS110. Interestingly, Cmv6323 and Cmv9109 are part of the two groups that triggered the most consistent and strong cell death responses in *N. benthamiana*. This similarity, however, was not observed in the amino acid pairwise alignment of these proteins, for which the pair Cmv5316 and Cmv5317 have a higher percentage identity than the Cmv6323 and Cmv9109 pair.

The two predicted disulphide bonds shared across all predicted structures are conserved with the two predicted disulphide bonds present in all members of the Ecp32 family from *D. septosporum* and *F. fulva* (except Ds/FfEcp32-5, which does not have any cysteine residues), which suggests this is a conserved feature in proteins belonging to this family (Figure 5) (Tarallo et al., 2022).

The structure of the truncated Ecp32 candidate Cms4060 was also predicted since the protein triggered weak cell death in the same manner as Cmv5316 (Figure 4). Cms4060 has 77 amino acids less than the 226 expected for Cmv5316. The alignment between the predicted structures of Cms4060 and Cmv5316 (Figure 5) highlights the N-terminal conservation previously observed in the sequence alignment (Additional File 9); the predicted structure of Cms4060 (coloured green in the structural alignment) aligns with the region of Cmv5316 coloured in black, including one conserved disulphide bond (S-S1). The grey colouring represents the C-terminal region of Cmv5316 that is not present in Cms4060. Similarities in the structures of truncated protein Cms4060 and full-length Cmv5316, with conservation of cell death activity, might provide insights into the regions of the proteins from the Ecp32 family that are responsible for plant receptor or target recognition.

## Discussion

*C. minus* is an important pathogen of pines worldwide. However, very little is known about the biology of this pathogen. At first, differences in ascospore length and characteristics of the fungus grown on agar plates indicated that there were two morphological types of *C. minus* in New Zealand, termed *C. minus* ‘simile’ and *C. minus* ‘verum’ (Dick et al., 2001). However, with the development of molecular tools to differentiate these morphotypes, another morphotype was suggested to be present in the New Zealand *C. minus* population (Hunter et al., 2016). Host genotype is known to strongly influence symptom expression of CNC (Ismael et al., 2020; Suontama et al., 2019). With climate change upon us and seasonal variation shifting, it becomes more important to understand the biology of these morphotypes, to enable mapping of likely distributions, understand virulence and pathogenicity and how they might behave under future climatic conditions.

Through morphological and molecular analysis, we have confirmed the presence of a third *C. minus* morphotype in New Zealand, here classified as ‘novus’. Growth temperature data suggested optimum growth is different for ‘verum’ compared to ‘simile’ and ‘novus’. This in turn indicated that sporulation and germination may occur at different temperatures, which could influence the geographical distribution of these morphotypes, highlighting the importance of comparative studies between them.

To help shed light on how genetic differences in morphotypes might affect *C. minus* virulence or pathogenicity, genomes of isolates from the three morphotypes of *C. minus* were sequenced. This enabled the prediction and comparison of CEs. The effector prediction pipeline generated a first list of possible effector proteins for *C. minus*. Because effectors are often described as SSCPs (Lo Presti et al., 2015), these features were used to identify *C. minus* CEs. Then, these SSCPs were classified as apoplastic or cytoplasmic effectors according to EffectorP, resulting in a set of CE proteins from the eight *C. minus* isolates. However, other proteins that were not classified as SSCPs are also potential effectors that should not be discarded for future analysis. Nevertheless, this initial classification was a good starting point to functionally characterize *C. minus* effectors, and it would be interesting to assess which of these trigger cell death responses, using the non-host ATTA approach described previously.

Some pathogens are known to secrete effectors that occur in families (Denton-Giles et al., 2020; Lo Presti et al., 2015; Rocafort et al., 2022). Following the observed similarity between DsEcp32-1 (part of the Ecp32 family from *D. septosporum*) and Cms835 (from isolate NZFS809), it was assessed to what extent the Ecp32 protein family was also present in this and the other seven isolates of *C. minus*. The results indicated that different morphotypes have different numbers of family members, with the morphotypes also showing differences in pseudogenisation of *Ecp32* genes. This was also observed in the Ecp32 family from *D. septosporum* and *D. pini* (both pine pathogens) and *F. fulva*, as they have four, three and five family members respectively, with the presence of pseudogenes in some (Tarallo et al., 2022).

The diversification of gene families can occur due to tandem and/or segmental duplications (Leister, 2004), which both seem to be present in the expansion of this particular effector family from *C. minus*. For example, the genes encoding proteins Cms4058, Cms4059 and Cms4060 (truncated), from NZFS809, are on the same scaffold and adjacent to one another. This is different from what was observed in *D. septosporum* and *F. fulva*, where the location of the *Ecp32* genes are conserved in matching chromosomes between the two species but none of these family members are localized on the same chromosome (Mesarich et al., 2023). The differences in the number of family members and pseudogenes in each morphotype might reflect pathogen virulence and host range, as the selection pressure imposed by host *R* genes on plant pathogenic fungi can lead to an expansion and sequence divergence of an effector gene family (Denton-Giles et al., 2020; Franceschetti et al., 2017; Pendleton et al., 2014).

Also consistent with the predicted structures of Ecp32 proteins from *D. septosporum* and *F. fulva*, predicted structures generated for representative members of the *C. minus* Ecp32 family were shown to adopt a β-trefoil fold. This is a fold present throughout different kingdoms that is involved in many different processes and interactions between organisms (Žurga et al., 2015). The most common types of proteins that share this fold are trypsin inhibitors and lectins, many being virulence factors in plant-pathogenic fungi (Juillot et al., 2016; Schubert et al., 2012; Varrot et al., 2013). It remains to be determined whether these β-trefoil proteins from *C. minus* have any virulence functions when infecting the host. It would be interesting to determine the structures of other CEs from *C. minus* and identify to what extent this fold is present in the structures of any of those proteins. For example, the Ecp20 family, another cell death-eliciting family present in both *D. septosporum* and

*F. fulva* (Tarallo et al., 2022), could not be detected in *C. minus* through primary sequence similarity searches.

It remains possible, however, that searches in the predicted structures of all *C. minus* CEs might identify proteins that adopt the characteristic Alt a 1 fold of Ecp20 family proteins, thereby expanding the repertoire of common effector types between these foliar fungal pathogens.

Proteins from the Ecp32 family in *C. minus* were found to have members that differed in their ability to trigger cell death in the non-host species *N. benthamiana* and *N. tabacum*. This difference was also observed in members of the Ecp32 family from *D. septosporum* and *F. fulva* (Tarallo et al., 2022). Whether the identified sequence variation contributes to the virulence functions of these proteins, as well as cell death elicitation, remains to be determined. Of note, other experiments, such as reactive oxygen species measurements and an analysis of defence marker gene expression in *Nicotiana* species, should now be performed to determine if these cell death responses are actually due to activation of the plant immune system. Similarly, the CE proteins should be transiently expressed in *BAK1*/*SOBIR1*-silenced or knock-out plants of *Nicotiana* species, to determine whether BAK1 and SOBIR1 (extracellular co-receptors involved in transducing defence response signals following apoplastic effector recognition) (Wang et al., 2018) are required for the elicitation of cell death responses by these Ecp32 family members. The differences in cell death-triggering activity observed between different family members might be due to differences in the composition and positioning of amino acids on the surface on the proteins, for example as a result of insertions, deletions and/or substitutions of residues, which occur more often in loops than in α-helices and β-sheets (Fiser et al., 2000). In some cases, loops in tertiary structures of proteins are known to be involved in protein-protein interactions, recognition sites, signalling cascades and ligand binding (Fetrow, 1995), all of which could explain why some proteins from the *C. minus* Ecp32 family trigger more consistent cell death responses than others. Also, sequence and structural diversification observed for the proteins in this family could have a role in evasion of host immunity, another possible explanation for why this family is expanded in *C. minus*.

The finding that Cms4060 (truncated) and Cmv5316 (full-length) triggered the same degree of cell death in both *Nicotiana* species suggests that the N-terminal part of the protein (that is also conserved between Cms4060 and the other Ecp32 proteins) might contain the region, or conserved epitope, that is either recognized by a plant extracellular immune receptor or is interacting with the plasma membrane, triggering cell death. Previous studies have demonstrated the possibility of swapping loop regions from non-functional proteins for other loops from functional proteins (Pardon et al., 1995; Wolfson et al., 1991). Considering the consistent cell death triggered by Cmv9109, Cms9849 and Cmn3003, it would be informative to try to determine differences in the loop regions of these proteins that might be responsible for cell death, in comparison with other proteins that trigger inconsistent or no cell death responses. It would then be possible to determine if there are differences between the *C. minus* morphotypes in how these proteins function.

In conclusion, the genome sequences of eight *C. minus* isolates supported taxonomic classification of *C. minus* ‘novus’ as a third morphotype in the New Zealand population of this pathogen. It is important to understand the geographical range and variations in virulence and/or pathogenicity of these morphotypes, as well as the relationships between host genotype and fungal morphotypes, all of which have implications for diagnostics and biosecurity for the forest industry. It was also possible to identify CE proteins that might be good candidates for future investigation in the *C. minus*-pine interaction. We also showed that, just as in *D. septosporum* (another pine pathogen) and *F. fulva*, the Ecp32 family is present in *C. minus*, and the evidence suggests that some members from this family might be recognized by plant immune receptors. This, combined with the fact that these proteins are conserved at the primary sequence and tertiary structure levels, suggest that they have important and/or conserved roles in fungal virulence. Amino acid sequence searches revealed that the Ecp32 protein family is widespread across other fungal species (Tarallo et al., 2022) and, therefore, might be an important core effector of plant-pathogenic fungi. Disease resistance based on core effectors that are vital for a pathogen’s ability to cause disease is more likely to be durable. Knowledge of core effectors from different *C. minus* morphotypes can enable the development of molecular tools that can complement breeding programmes, not only in this, but also in other forest pathosystems.

## Data availability

The sequences were deposited in NCBI under the accession numbers outlined in Table 2.

## Conflict of interest

The authors declare that the research was conducted in the absence of any commercial or financial relationships that could be construed as a potential conflict of interest.

## Authors contributions

MT completed the effector analysis and wrote the manuscript. KD performed the culture-based analysis (with assistance from KG) and contributed substantially to the manuscript. LNL, TW and DS performed the DNA extraction and genome sequencing analysis. RM, RB and CM conceived and supervised the study, made substantial contributions to the interpretation of the results, and contributed to writing and revision of the manuscript. All authors contributed to the article and approved the submitted version.

## Funding

This research was funded by Scion (New Zealand Forest Research Institute, Ltd., Rotorua, New Zealand) through the Resilient Forests Research Program via Strategic Science Investment Funding from the New Zealand Ministry of Business Innovation and Employment (MBIE), grant number CO4X1703, and New Zealand Forest Grower Levy Trust funding, grant number QT-9108.

## Acknowledgements

The authors thank Rita Tetenburg and Sara Carey for technical assistance with culture collections and Dale Corbett for help with graphic design (Scion, Rotorua, New Zealand). All strains used in this study are housed and maintained in Scion’s Forest Health Reference Laboratory Culture Collection.

## Additional Files Legends

**Additional File 1.** Accessions for phylogenetic tree used in Additional File 2.

**Additional File 2.** Multi-locus phylogeny of *Cyclaneusma* species and other fungal associates of *Pinus radiata* showing the placement of the three *Cyclaneusma* morphotypes. Complete or partial DNA sequences from three gene loci (Internal Transcribed Spacer [ITS] of ribosomal DNA, nuclear LSU [nLSU] and translation elongation factor 1α [tef-1]) were used to infer phylogeny using Maximum Likelihood in Geneious Pro v10.0 and Mega X. Scale bar shows number of substitutions per site. *Gremmeniella abietina* DAOM 170367 was used as an outgroup (available at https://mycocosm.jgi.doe.gov/Greabi1/Greabi1.home.html).

**Additional File 3.** Descriptions of colony morphology of ‘verum’, ‘smile’ and ‘novus’ morphotypes on 2% and 9% MEA at 22°C in the dark after 8 weeks.

**Additional File 4.** Temperature relationships between *Cyclaneusma minus* morphotypes, ‘simile’ (blue), ‘verum’ (red) and ‘novus’ (green), using mean radial growth, measured in millimetres per day, with standard deviations.

**Additional File 5.** *Cyclaneusma minus* isolates genome sequencing results.

**Additional File 6.** Protein sequence for the predicted candidate effectors from *Cyclaneusma minus* isolates NZFS110, 725, 1800, 3276 (morphotype ‘verum’), NZFS759, 809 (morphotype ‘simile’) and NZFS3305, 3325 (morphotype ‘novus’).

**Additional File 7.** Amino acid sequences of *Cyclaneusma minus* Ecp32 family members in eight isolates.

**Additional File 8.** Protein phylogeny of the Ecp32 families from *Dothistroma septosporum*, *Fulvia fulva* and *Cyclaneusma minus* NZFS110. The tree was constructed with the neighbour-joining method using Geneious Software v9.1.8. Bootstrap values are shown at the nodes as percentages. The scale bar represents 0.3 substitutions per site. Ds: *D. septosporum*; Ff: *F. fulva*; Cmv: *C. minus* morphotype ‘verum’.

**Additional File 9.** Protein sequence alignment of Ecp32 family members across eight isolates and three morphotypes of *Cyclaneusma minus*. The alignment was generated using Clustal Ω (Sievers et al., 2011), with blue shading indicating the degree of amino acid sequence conservation. Conserved cysteine residues are highlighted in black and predicted signal peptides are underlined. Intrinsically disordered regions predicted with Predictor of Natural Disordered Regions (PONDR^®^ VLXT) (Romero et al., 2001) are underlined in red. The number following each morphotype ID (NZFS) refers to the protein number from the annotated genomes; the asterisk * indicates that the protein was incorrectly predicted, and the corrected protein sequence was used. Cms: *C. minus* morphotype ‘simile’; Cmv: *C. minus* morphotype ‘verum’; Cmn: *C. minus* morphotype ‘novus’. Proteins were colour-coded according to their clustering in the phylogeny in Figure 3.

**Additional File 10.** Ecp32 proteins from different morphotypes of *Cyclaneusma minus* trigger cell death in *Nicotiana tabacum*. (A) Proteins were expressed in *N. tabacum* using an *Agrobacterium tumefaciens*-mediated transient expression assay to assess their ability to elicit cell death. Representative images are shown (n = 12– 30 infiltration zones), from at least three independent experiments. INF1, *Phytophthora infestans* elicitin positive cell death control; EV, empty vector negative no-cell death control; Cms: *C. minus* morphotype ‘simile’; Cmv: *C. minus* morphotype ‘verum’; Cmn: *C. minus* morphotype ‘novus’. Photos were taken 7 days after infiltration. (B) Graphs display the percentages of infiltration zones that showed cell death in response to each different protein, divided into four categories: strong cell death, weak cell death, chlorosis, and no cell death. The coloured line under the protein ID refers to the different gene and protein groups in Figures 2 and 3.

**Additional File 11.** Predicted local-distance difference test (pLDDT) score and global superposition metric template modelling score (TM-score) for *Cyclaneusma minus* Ecp32 proteins structural predictions.

## Notes

### Competing Interest Statement

The authors have declared no competing interest.

