## Supplementary figures and images for "Genomic, effector protein and culture-based analysis of *Cyclaneusma minus* in New Zealand provides evidence for multiple morphotypes"

### Additional File 2

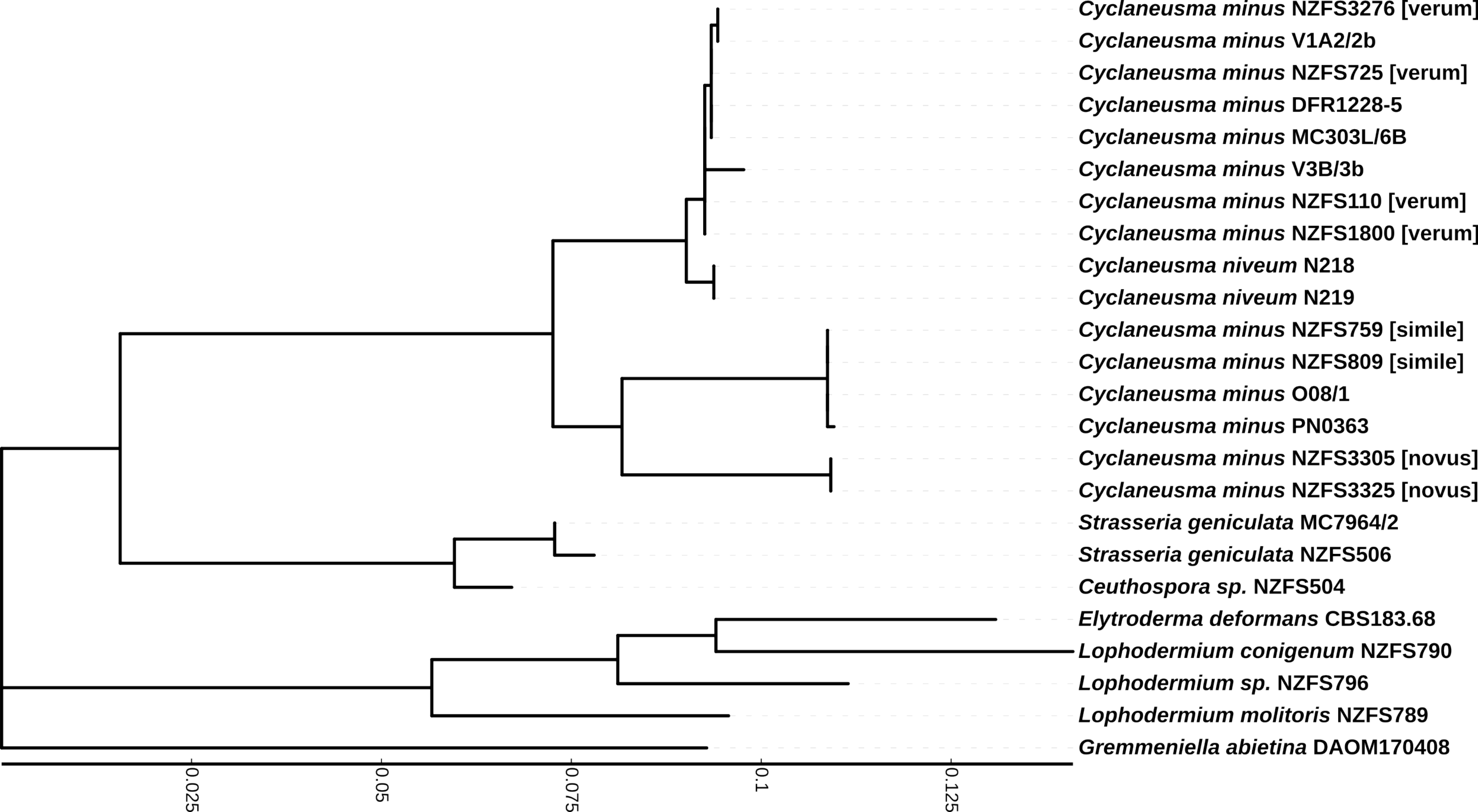

### Additional File 4

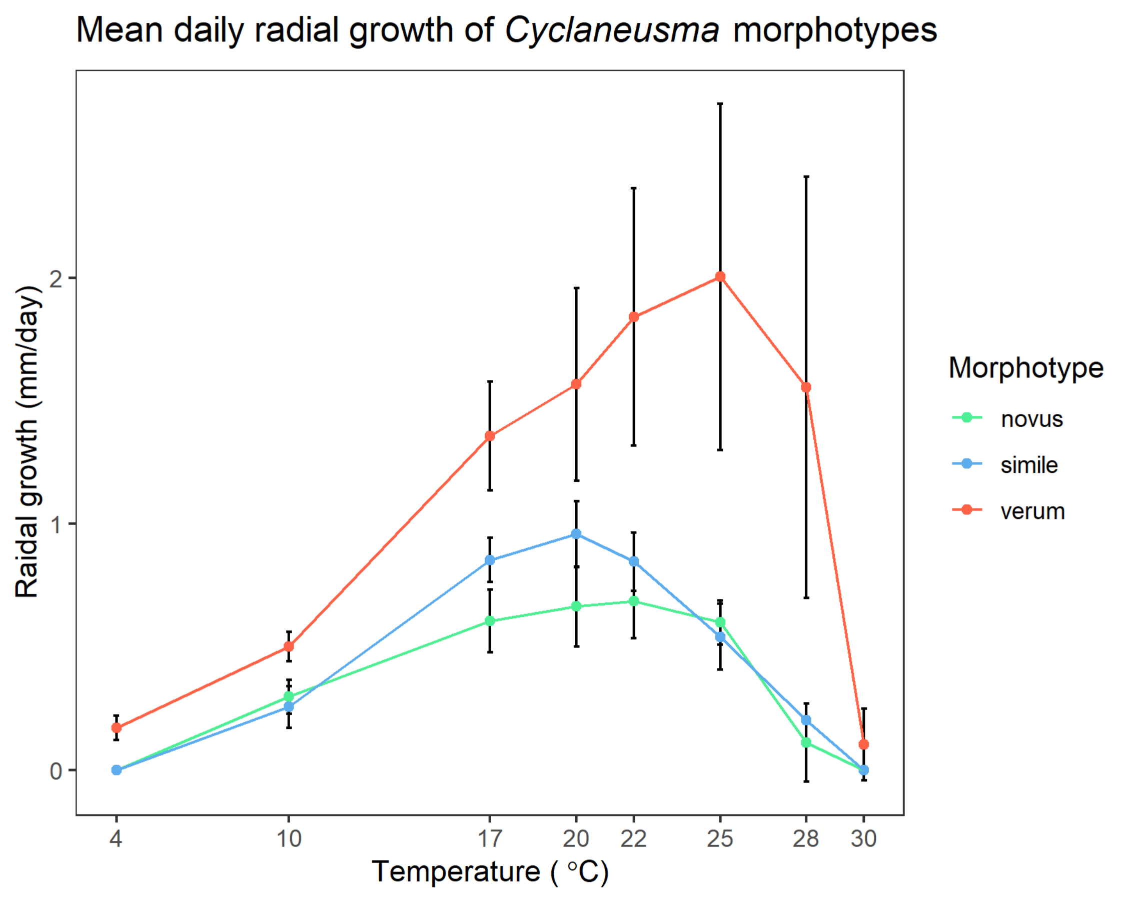

### Additional File 8

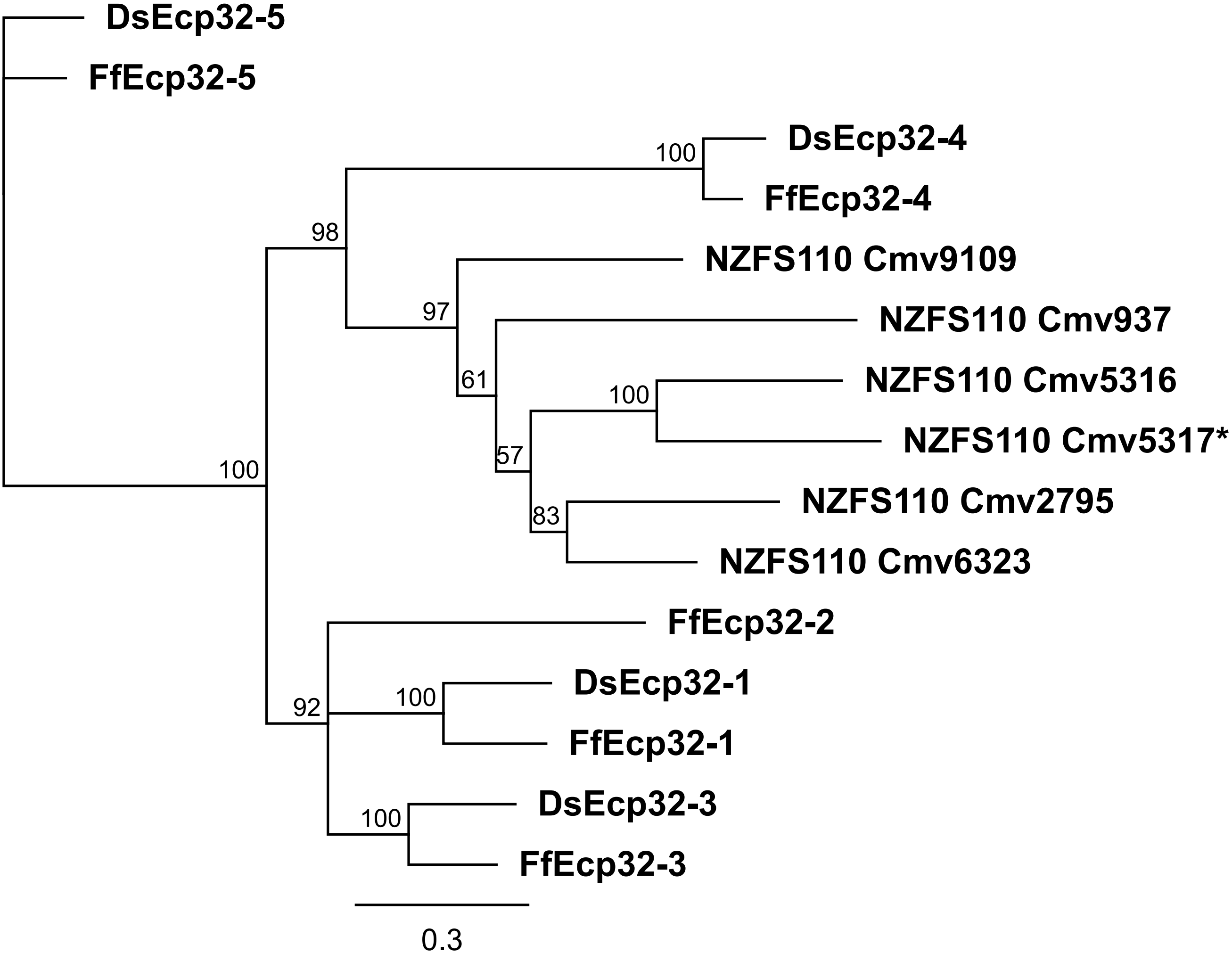

### Additional File 10

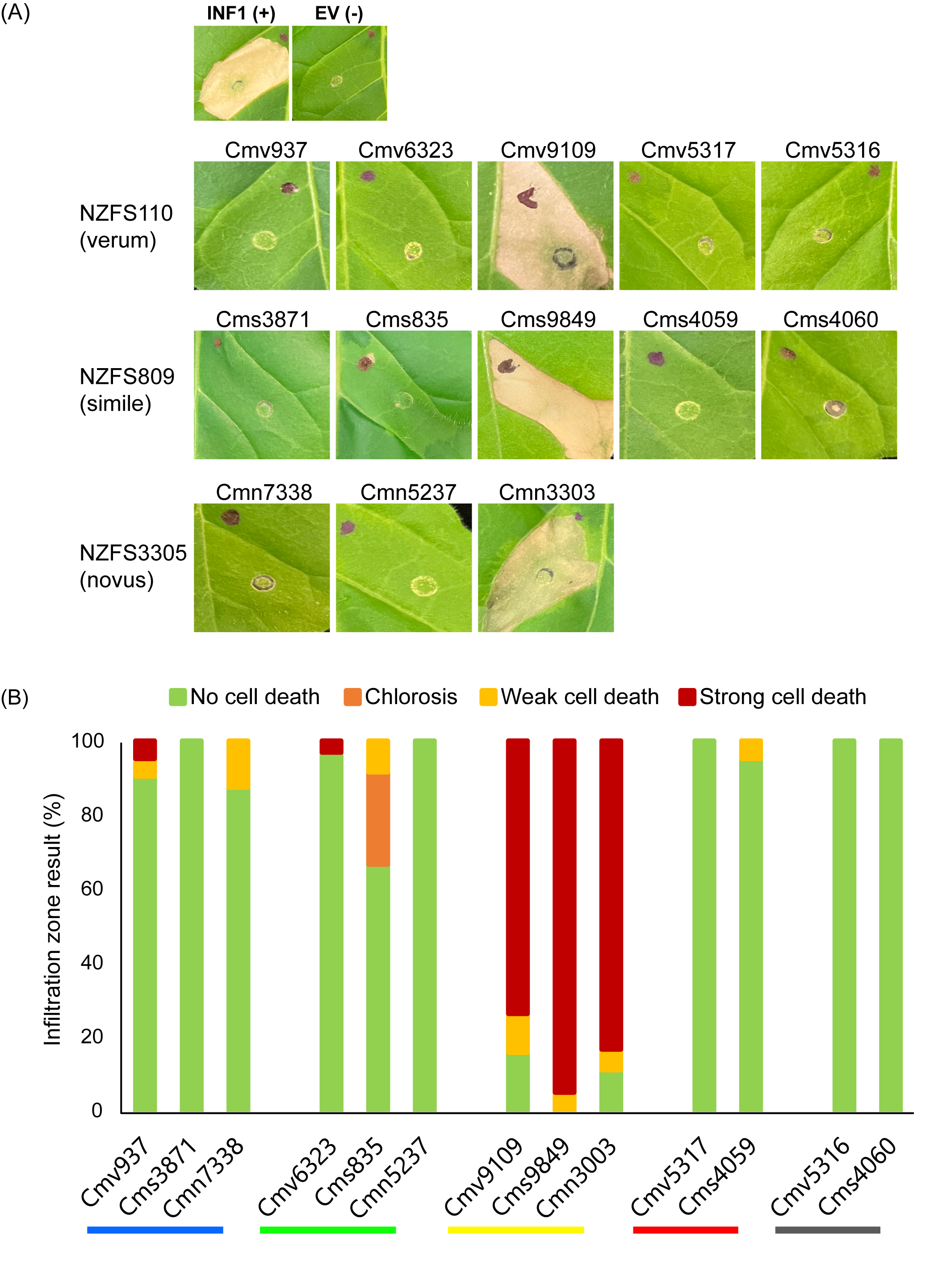
