## Additional File 9 for "Genomic, effector protein and culture-based analysis of *Cyclaneusma minus* in New Zealand provides evidence for multiple morphotypes"

```
NZFS809 Cms4060        1 -MFAITVLVGCLSAL---TAASPMLTARDH-----------------------------------------------   26
NZFS110 Cmv5317*       1 -MSAAIFLLAFISSL---CSASPL--TTLS-----------------------------------------------   24
NZFS809 Cms4059        1 -MITTTILITLLSTL---TRSSLL-QNKRQ-----------------------------------------------   25
NZFS110 Cmv5316        1 -MCAATLLMGFLSAL---AAASPLIQAQRR-----------------------------------------------   26
NZFS725 Cmv1064        1 -MCAATLLMGFLSAL---AAASPLIQAQRR-----------------------------------------------   26
NZFS3276 Cmv2969       1 -MCAATLLMGFLSAL---AAASPLIQAQRR-----------------------------------------------   26
NZFS110 Cmv937         1 -MLSATLLAVLLPAL---TCASPLNFLLPRQND--------------------------------------------   29
NZFS809 Cms3871        1 -MLSTTLLAAFLPAL---TLASSLSSLLPRQD---------------------------------------------   28
NZFS3305 Cmn7338       1 -MLAPTVLAALLPIL---TSASPLDFLVPRQGTGA------------------------------------------   31
NZFS809 Cms4058        1 -MFTAAISLTLLSALFILSKASPLT---TR-----------------------------------------------   26
NZFS110 Cmv9109        1 -MHAVAAIVAILPAL---AAASPLA---KRA----------------------------------------------   24
NZFS3276 Cmv4749       1 -MHAVAAIVAILPAL---AAASPLA---KRA----------------------------------------------   24
NZFS809 Cms9849        1 --MYAAALAAFLPAL---AAASPLA---KRN----------------------------------------------   23
NZFS3305 Cmn3003       1 -MYAAIIAAAVLPAL---AVAGPIA---ERA----------------------------------------------   24
NZFS110 Cmv2795        1 MLFTPKLLAVLLPAL---AAASTIP---KRTDSGMSGDKSAAMSPETFGAAPVPVLGDKTPDNSGDKYPVSGAGSSN   71
NZFS3276 Cmv3467       1 MLFTPKLLAVLLPAL---AAASTIP---KRTDSGMSGDKSAAMSPDTFGAAPVPVLGDKTPDNSGDKYPVSGAGSSN   71
NZFS110 Cmv6323        1 MVATAAVLATLLPAL---AAASPLK---QR-----------------------------------------------   24
NZFS3276 Cmv3921       1 MVATAAVLATLLPAL---AAASPLK---QR-----------------------------------------------   24
NZFS809 Cms835         1 -MFTPTLLTALLPAL---AAASPIF---KR-----------------------------------------------   23
NZFS3305 Cmn5237       1 -MFATTLLAAVLPAL---AAASPIK---RD-----------------------------------------------   23

NZFS809 Cms4060       27 -------------YQNPFETGMIVGGPSNSPIYFDSISAANSSFWIGADTRASCNSS-----ISCSIYPNDTSIIQY   85
NZFS110 Cmv5317*      25 -------------PPPFLATSLVAIGPLNTPIYTQPIVANSTFFWIGTTTRTSCPHFS--NNFTCAIYPNDTSLLIY   86
NZFS809 Cms4059       26 -------------ETSFLATGLTAIGNLNTPVYGQTLNANSSHFWLHTTTRTSCPTFS--NNFTCAIYPNDTSILVY   87
NZFS110 Cmv5316       27 -------------ETPYLASSFISVGTPNTPIYFDTISAANSSFWIGASTRTYCPSSSPENFTNCSAFQNSTSILVY   90
NZFS725 Cmv1064       27 -------------ETPYLASSFISVGTPNTPIYFDTISAANSSFWIGASTRTYCPSSSPENFTNCSAFHNSTSILVY   90
NZFS3276 Cmv2969      27 -------------ETPYLASSFISVGTPNTPIYFDTISAANSSFWIGASTRTYCPSSSPENFTNCSAFHNSTSILVY   90
NZFS110 Cmv937        30 -------------THPDESAGMIALASGT-VIHFGTVNASDNAFYIGRPTDTYCPSTP----DICGGLLNITMIHPY   88
NZFS809 Cms3871       29 -------------SHQSENAGMIALHSGS-VIHFSTINASDSIFYIGIPTGTYCPDTV----PGCSGLTNTTVIHPY   87
NZFS3305 Cmn7338      32 -------------THSNQPAALAALDSGS-PIHYQSINASDNQFYIGLPTDVYCPLEP----PLCAGFPNITAVYPD   90
NZFS809 Cms4058       27 -------------QDGYYTTSLLSIGRTPSWIFFEPVNATSSLFWIGAPTDAACPT-P----YDCSGFSNTTVIRVD   85
NZFS110 Cmv9109       25 -------------ITFPYSAEWIALRSAS-PVHFQSLNANSSTIWIGKETATYCDP-V----ADCESYINTTSITL-   81
NZFS3276 Cmv4749      25 -------------ITFPYSAEWTALRSAS-PVHFQSLNANSSTIWIGKETATYCDP-V----ADCESYINTTSITL-   81
NZFS809 Cms9849       24 -------------ITFPYSAEWTALRSGSEYIHYQSLNANSQLIWIGKETNVDC-P-I----GGCESYSNTTSITL-   80
NZFS3305 Cmn3003      25 -------------ITFPYSATLTSLRSGS-DIQYAAINANSQAFWIGKETDYDC-P-I----GGCESYSNTTSITI-   80
NZFS110 Cmv2795       72 IFGAAPVPVSGDKSDTPYTTGLVAIRSGS-DFQYASVNASSGLFYIGESTKTGCAA-V----ANCGNYPNITLITIH  142
NZFS3276 Cmv3467      72 IFGAAPVPVSGDKSDTPYTTGLVAIRSGS-DFQYASVNASSGLFYIGESTKTGCAA-V----ANCGNYPNITLITIH  142
NZFS110 Cmv6323       25 ----------AFLADYPYTTGLEALRSAS-PIHFSAVNASLGHFFIGKATNTYCP--V----SDCSGFPNITVISIY   84
NZFS3276 Cmv3921      25 ----------AFLADYPYTTGLEALRSAS-PIHFSAVNASLGHFFIGKATNTYCP--V----SDCSGFPNITVISIY   84
NZFS809 Cms835        24 ----------DDY-VPPYTTSLLSLRSAS-PIHFSAINASHGHFSIGEPTSYYCP--V----SDCSGFPNITLISIY   82
NZFS3305 Cmn5237      24 ----------V---EYPYTTGLEALRSTS-PIHFQAVNASLGQFWIFKDTDTYCP--I----SSCASYPNITSINIY   80

NZFS809 Cms4060       86 E--D--VAWMTTAVAGSQNVYVAESVALSFTEPHASFVNNTYPPGSYIP--EFLIW-----GRPELVFVLIVLG-R*  150
NZFS110 Cmv5317*      87 P--S-SSAYMNSAVPGSQNLYVAESGALQFTEPHAE---GAYPAGSYTS--GFNVE---MYGAANIDPTVNFTG-GG  151
NZFS809 Cms4059       88 N--SSSTAYMDSSVPGSQNIFVAPSGALSFTEPHAE---GIFPEGSYVS--GFQMV---EYGH-GIDPLLNFTG-GG  152
NZFS110 Cmv5316       91 N--TSTTASMNSAVPGGQTIYVSPTGALSFTSPNR----SIIPAGSYTS--GFTIGCEQPYYPPECSPTWNFTG-GW  158
NZFS725 Cmv1064       91 N--TSTTASMNSAVPGGQTIYVSPTGALSFTSPNR----SIIPAGSYTS--GFTIGCEQPYYPPECSPTWNFTG-GW  158
NZFS3276 Cmv2969      91 N--TSTTASMNSAVPGGQTIYVSPTGALSFTSPNR----SIIPAGSYTS--GFTIGCEQPYYPPECSPTWNFTG-GW  158
NZFS110 Cmv937        89 DT--G---VINMQAIQEQMVFVDATGALSFTTAHEEDIT---PPGSYVS-GGFMVE---E--ESS-VSLLQFDQ-GA  149
NZFS809 Cms3871       88 NP-DG---KVYMGSINVQQLYVEPTGALAFSTVHEEDIF---PPGAYTSSGDFTVD---E--ESS-VSLLEFGA-GS  150
NZFS3305 Cmn7338      91 DG-GRDGAGMDAVAPGGQDVFVAESGALMFTTAHEEDDY---PPNSYTD-GGFEVQ---EYEDGA-LSELAFTA-GD  157
NZFS809 Cms4058       86 RDPSQASAFMDSPVAGGQMLYVTESGALSFMQPHCGSCGI--TNADSLT--GFGIN---YNGNGTGSDLLSFEVDRV  155
NZFS110 Cmv9109       82 ---RSGNAYMNVDVAGGQQLYIAASGAIEFTQPHSAA----MPEGSVTG--GWGAY---TSENGI--DMITNSAG--  142
NZFS3276 Cmv4749      82 ---RSGNAYMNVDVAGGQQLYIAASGAIEFTQPHSAA----MPEGSVTG--GWGAY---TSENGI--DMITNSAG--  142
NZFS809 Cms9849       81 ---SSGNAYMNDVVAGGQQLFVAADGALEFTVVHSAA----IPEGAFVG--GWGAY---TTENGV--DMITFSSGGD  143
NZFS3305 Cmn3003      81 ---SGQSATMNSVVAGGQAVYVTLSGAIEFTQAHSAA----KPEGAITG--GWGAS---TTANGV--DIITFTGGG-  142
NZFS110 Cmv2795      143 E--TEGTAYLDADVPAGQQVFVAQSGALSFARPHQQSAFVSSPHGPYSN--GFGVY---NKGDGASFDILNFTLA--  210
NZFS3276 Cmv3467     143 E--TEGTAYLDADVPAGQQVFVAQSGALSFARPHQQSAFVSSPHGPYSN--GFGVY---NKGDGASFDILNFTLA--  210
NZFS110 Cmv6323       85 E--ASGTAYMSSNVPGGQNVFVAKSGALSFTEPHQEYVF---PDGSYTR--GFEIA---NNGHGV--DLLNFTGGG-  148
NZFS3276 Cmv3921      85 E--ASGTAYMSSNVPGGQNVFVAKSGALSFTEPHQEYVF---PDGSYTR--GFEIA---NNGHGV--DLLNFTGGG-  148
NZFS809 Cms835        83 E--ASGTAYMSTNVPGGQNVYVAESGALSFTEPHQEYVF---PDGSYTT--GFAVS---NNGNGV--DLLNFTQA--  145
NZFS3305 Cmn5237      81 E--SSGTASMSTDVPGGQDVFVAESGALSFTEPHQEFSF---PDGAYTT--GFAVT---NNGNGV--DLLNFTEA--  143

NZFS809 Cms4060          -------------------------------------------------------------------------
NZFS110 Cmv5317*     152 ATAFWACPPNDPAENSNPEGTGIDDPYKIFLDVEGWNDSVVPSGNISHCLPFNIATKQDAYGDGEVHGAYQYI*  225
NZFS809 Cms4059      153 ATAFWGCPMSSEY--------VEDAPYQVFLNVEGWNDGVVPSGNTSDCIPFGVATNQDAYDDGEVHGAYQYI*  218
NZFS110 Cmv5316      159 ATGFYACPLNDPAVYADPY--GRDFPYKVFLDVEGW---VVPSGNMSDCLPFSVAVSPDMYSDGVAWGAYEYV*  227
NZFS725 Cmv1064      159 ATGFYACPLNDPAVYADPY--GRDFPYKVFLDVEGW---VVPSGNMSDCLPFSVAVSPDMYSDGVAWGAYEYV*  227
NZFS3276 Cmv2969     159 ATGFYACPLNDPAVYADPY--GRDFPYKVFLDVEGW---VVPSGNMSDCLPFSVAVSPDMYGDGVAWGAYEYV*  227
NZFS110 Cmv937       150 SQGWVACPITSDVDENG----QPLGPFQVFASVVGFSDLVAPLGCTEDCIGFDMALE--WISAASPPGAYEYD*  217
NZFS809 Cms3871      151 SQGWVACAITNDLDESG----QLLGPWQVFAEVEGFGDSVVPSGCTEDCIGIDIALA--WESVTVPPSAYAYD*  218
NZFS3305 Cmn7338     158 ASSWLACPITDDVDDSG----MQLGPWQVFADVVGFNDSVVPLGCVDECLGFDMLLQ--WIDVDAVEAAFEYV*  225
NZFS809 Cms4058      156 AKGFFACPPTHINVTNG----NEEFPYQVFLDVKGWNDSVVPSGRTVDCFGFEIAVH--DMGSDAMEGASEYV*  223
NZFS110 Cmv9109      143 -TGFLACPT-----TD------L-TSYQVFADIEGIPNSAVPSGNTDDCLGFDIAAS--VV-NDA--TAYEYL*  198
NZFS3276 Cmv4749     143 -TGFLACPT-----TD------L-TSYQVFADIEGIPNSAVPSGNTDDCLGFDIAAS--VV-NDA--TAYEYL*  198
NZFS809 Cms9849      144 QGGFLACPM-----KD------SDEVYQIFADIEGIPNSAVPSGNTADCLGFDIAAS--KV-DSP--TAYQYD*  201
NZFS3305 Cmn3003     143 ATTFLACPA-----ST------GGSPYQVFANVEGMVNSGVPSGNVADCLDISIAGS--VV-EGA--TAYQYD*  200
NZFS110 Cmv2795      211 -NTFQACPSS----KK------GGKPYQVFLDVPGFKDHVVPSGKVADCLPFEIAVS--SYKTGA--GAYQYE*  269
NZFS3276 Cmv3467     211 -NTFQACPSS----KK------GGKPYQVFLDVPGFKDHVVPSGKVADCLPFEIAVS--SYKTGA--GAYQYE*  269
NZFS110 Cmv6323      149 ATTFTACPPTTPSSTA------GVYPYQVFLDVKGFNDSVVPSGSVADCLPFGIAIY--NS-TGA--GAFQYE*  211
NZFS3276 Cmv3921     149 ATTFTACPPTTPSSTA------GVYPYQVFLDVKGFNDSVVPSGSIADCLPFGIAIY--NS-TGA--GAFQYE*  211
NZFS809 Cms835       146 -NTFTACPPC----TG------GEFPYQVFLDVPGFNDSVVPSGSTADCLPFGIAIN--NYTTDA--GAFQYE*  204
NZFS3305 Cmn5237     144 -ESFTACPV-----TP------GEAPYQVFLTVPGFNDSVVPSGSTADCLGFGIGIY--NS-TGK--GAFQYE*  200
```
